# Piriform cortex provides a dominant gamma LFP oscillation in the anterior limbic system

**DOI:** 10.1101/861021

**Authors:** James E. Carmichael, Matthew M. Yuen, Matthijs A. A. van der Meer

**Affiliations:** Department of Psychological and Brain Sciences, Dartmouth College, Hanover NH 03755

**Author notes:** Correspondence should be addressed to MvdM, Department of Psychological and Brain Sciences, Dartmouth College, 3 Maynard St, Hanover, NH 03755. **Conflict of Interest**: The authors declare no competing financial interests.

## Abstract

Oscillations in the local field potential (LFP) are widespread throughout the rodent limbic system, including in structures such as the orbitofrontal cortex and nucleus accumbens. Synchrony between LFPs across these structures, as seen during specific behavioral events, is often interpreted as evidence of a functional interaction. However, the source of these oscillations is often tacitly assumed to be local, leading to a potential misattribution of function. Using *in vivo* simultaneous multisite recordings in freely moving male rats (n = 7) we demonstrate that gamma-band LFP oscillations (45-90 Hz) in multiple anterior limbic structures are highly synchronous not only with each other, but also with those in piriform cortex. Phase reversals across the piriform cortex cell layer and susceptibility to nasal occlusion indicate that piriform cortex is the source of these common gamma oscillations. Thus, gamma-band LFP oscillations seen in brain regions adjacent to the piriform cortex are likely not generated locally, but are instead volume conducted from the piriform cortex. This emerging view of gamma oscillations in anterior limbic circuits highlights the importance of the common piriform cortex input as a major influence and introduces caveats in the interpretation of locally recorded LFPs.

## Introduction

Many aspects of brain function are thought to benefit from rapid changes in the ability of one population of neurons to affect another. A prominent proposal of how such dynamic connectivity can be accomplished is through the temporal coordination of neural activity (synchrony). Local field potentials (LFPs), which reflect the summed transmembrane currents within an area, offer a powerful experimental access point for probing circuit- and systems-level communication. Specifically, oscillations visible in the LFP have been implicated in routing the flow of information between brain structures (Colgin et al., 2009; Bosman et al., 2012; Igarashi et al., 2014) by creating temporal “windows of opportunity” during which postsynaptic excitability is highest (Fries, 2005; Colgin, 2013; Womelsdorf et al., 2014; Fries, 2015; Bonnefond et al., 2017). Although the mechanistic relevance of oscillations in brain activity remains an active area of debate, in any case LFP oscillations can provide a useful readout of different behavioral, cognitive (Jones and Wilson, 2005; Gruber et al., 2009; Shin et al., 2017; Igarashi et al., 2014; Wu et al., 2018), and clinical brain states (Brown et al., 2001; Cassidy et al., 2002; Uhlhaas and Singer, 2006; Jadi et al., 2016).

Limbic structures such as the prefrontal cortex (PFC), hippocampus (HC), amygdala (Amg), and nucleus accumbens (NAc) interact to support a variety of behaviors including goal-directed decision making, navigation, and memory recall. Rich patterns of LFP oscillations have been recorded within each of these sites individually, and LFP synchrony across multiple sites has been associated with conditioned freezing and anxiety (Adhikari et al., 2010b; Likhtik et al., 2014; Karalis et al., 2016; Moberly et al., 2018; Concina et al., 2018), working memory (Jones and Wilson, 2005; DeCoteau et al., 2007; Fujisawa and Buzsáki, 2011; Place et al., 2016), and rodent models of psychiatric conditions (Sigurdsson et al., 2010; Stujenske et al., 2014). Given the utility of these limbic LFP patterns as a biomarker and measure of functional connectivity (Harris and Gordon, 2015), and the possibility of mechanistic relevance, it is important to identify their source(s) in the brain, which is often tacitly assumed to be local (i.e. generated where the LFP is recorded).

However, an emerging body of work has identified non-local source(s) for prominent oscillations in limbic system LFPs across a range of frequency bands. For example, theta oscillations (7-10 Hz) have been recorded in multiple regions such as the dorsal striatum (dStr) and PFC. But for both the dStr (Lalla et al., 2017) and the PFC (Sirota et al., 2008), the HC has been identified as the source of these theta oscillations. Similarly, gamma band oscillations (45-90 Hz) in the NAc have no local sources, but are instead volume-conducted from the adjacent piriform cortex (PC) (Neville, 2003; Berke, 2009; Carmichael et al., 2017). In an extreme example, whisker-evoked deflections in the mouse olfactory bulb (OB) LFP can travel via volume conduction to the orbitofrontal cortex (OFC) even when the connections between the two areas are severed (Parabucki and Lampl, 2017). Thus, not only can the source(s) of LFPs be non-local, the volume-conducted signal can travel long distances (see also Logothetis et al. 2007; Kajikawa and Schroeder 2011).

These examples of non-local oscillations permeating into other regions highlight the importance of determining the correct source of an LFP. First, without knowing the source, underlying activity and models of function can be misattributed to the wrong brain region. Second, knowing the source and mechanisms that generate the LFP can also lead to more effective targeting in clinical interventions. For instance, a change in NAc gamma oscillations could correlate with a pathological marker and thus become a target for deep brain stimulation (DBS), when the signal is actually reflecting PC activity. Given the importance of identifying the source(s) of the LFP, we sought to determine how widespread non-local LFPs are in the rodent limbic system. The PC in particular lies proximal to several brain regions with prominent gamma oscillations with various behavioral correlates. Thus, we expect that these piriform-proximal regions could also contain volume-conducted PC gamma, as is the case with the NAc.

Specifically, we recorded simultaneous LFPs during wakeful rest from electrodes in the prelimbic (PL), infralimbic (IL), cingulate (CG), piriform (PC), and orbitofrontal (OFC) cortices, as well as the nucleus accumbens (NAc). LFP gamma oscillations in these regions are correlated with different aspects of behavior, and local spiking activity shows phase-locking to these oscillations (rats: Berke 2009; van der Meer and Redish 2009; van Wingerden et al. 2010b; Kalenscher et al. 2010; Howe et al. 2011; Morra et al. 2012; Insel and Barnes 2015; mice: Sohal et al. 2009; Kim et al. 2016). Moreover, observations of highly synchronous gamma oscillations across these structures have been interpreted as evidence for a functional interaction between them (Dejean et al., 2013; Donnelly et al., 2014; Catanese et al., 2016). We address the possibility that this common gamma oscillation is volume-conducted from the PC into multiple neighboring limbic regions with three complementary approaches. First, we characterize the temporal relationships between gamma oscillations in each of these regions. If gamma oscillations are in fact volume-conducted from the PC, they would be highly correlated at fine timescales. Second, we employ a reversible nasal occlusion protocol (Kucharski and Hall, 1987; Cummings et al., 1997) known to abolish piriform (Zibrowski and Vanderwolf, 1997) and NAc (Carmichael et al., 2017) gamma power. Finally, we record from electrodes spanning the piriform cortex cell layers to determine if there is a phase reversal of gamma oscillations, which would pinpoint it as the source.

## Methods

### Overview

This study consists of two experiments: the *multi-site* and *trans-piriform* experiments. Both experiments followed the same overall procedures, but used different recording sites. The multi-site experiment targeted a set of recording locations across multiple limbic brain structures (described below), and the trans-piriform experiment focused on recording from sites on either side of the piriform cortex/olfactory tubercle pyramidal cell layer. For both experiments, local field potentials were acquired from male rats (n = 7 total across experiments) while they were awake and resting. Daily recording sessions consisted of four 10-minute recording blocks containing different experimental conditions to evaluate the effects of unilateral nostril blockage (naris occlusion, described in more detail below). All procedures were approved by the Dartmouth College IACUC (protocol # vand.ma.2).

### Subjects

In total seven male Long-Evans rats (>10 weeks old; >400 g at the time of surgery; 4 from Taconic Biosciences, 3 from Envigo) were used. Six of the subjects had previously been used to pilot behavior on an unrelated navigation task. Animals were implanted with recording probes (described below) and given >4 days to recover post-op. Electrodes were slowly lowered to their target areas over 4-7 days, after which the animals were food-restricted to 18 g/day and daily recording sessions commenced (4 total recording sessions per subject). Before each daily recording session *ad lib* access to water was removed for 8 hr, and resumed within 30 min of the end of the session. Subjects were kept on a 12 hr light/dark cycle, with all experiments performed during the light phase.

### Surgery and recording probes

Custom-designed microdrives (Grasshopper Machinewerks LLC) were used throughout these experiments. The “multi-site” drive had four independently movable tetrodes or stereotrodes targeting the prelimbic (PL) and/or infralimbic (IL) portions of the medial prefrontal cortex (mFPC), orbitofrontal cortex (OFC), nucleus accumbens (NAc), and cingulate gyrus (CG) (summarized in Table 1). Only histologically confirmed recording sites were included for analysis; note that in some animals, some intended sites were not confirmed so not every animal has every site (confirmed sites are shown in Figure 1A). In some subjects the mPFC tetrode was replaced with two vertically offset stereotrodes to record from the PL and IL simultaneously. The “trans-piriform” drives used the same basic layout, except that a vertically staggered tetrode-stereotrode pair (offset 1-1.5 mm) was used to allow for simultaneous recording above and below the piriform layer (Figure 1A). For both drive types, tetrodes and stereotrodes were gold-plated (Sifco 6355) to impedances between 300-500 kΩ (Nano-Z, White Matter LLC). Surgical procedures were as described previously (Malhotra et al., 2015). Briefly, the custom recording drives were implanted in the right hemisphere under isoflurane anaesthesia and secured to the skull with screws and C & B Metabond dental cement (Parkell). A skull screw above the cerebellum was connected to a pin on the microdrive and acted as both ground and a reference.

**Figure 1:**
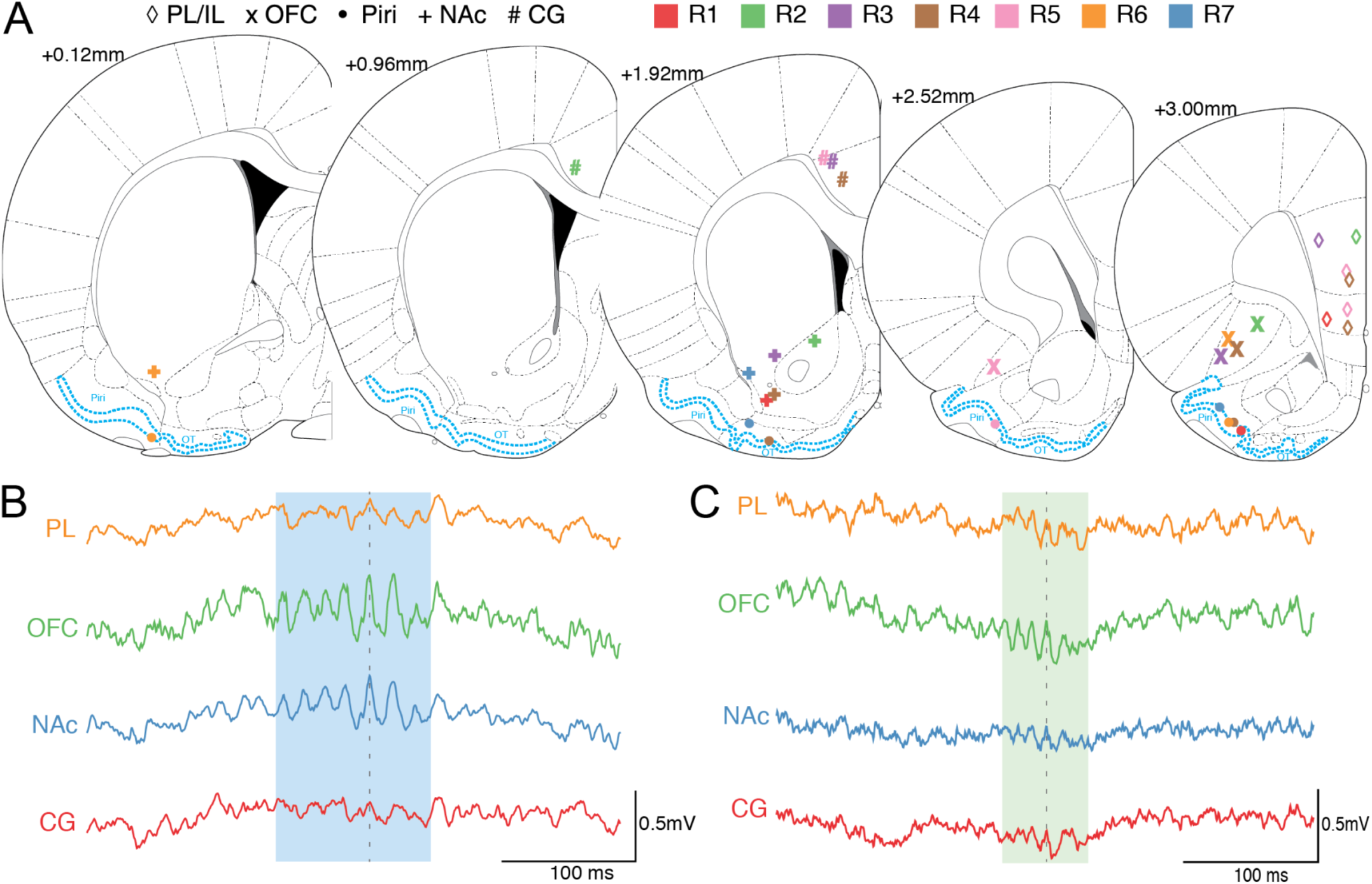
Gamma band oscillations in the LFP are prominent across multiple regions in the anterior limbic system. (**A**) Recording sites across all subjects for the structures of interest (PL: prelimbic; IL: infralimbic; OFC: orbitofrontal cortex; PC: piriform cortex; NAc: nucleus accumbens; CG: cingulate gyrus). The electrodes remained at a fixed depth across all recording sessions. Area highlighted in blue delineates the piriform cortex (Piri) and olfactory tubercule (OT); for brevity, we will refer to both these olfactory regions collectively as the piriform cortex. (**B**) Raw traces from four recording sites during a representative low-gamma event (blue shading) detected on the OFC channel. (**C**) Representative high-gamma event (green shading) in a separate subject from (**B**). Both the OFC and NAc show a strong oscillation with a similar number of cycles and little to no phase offset (grey dash line). The PL displays a more subtle oscillation at the same time as the OFC and NAc, while the CG is less clear but still present.

**Table 1:** Electrode targets used when implanting the multi-site drives. Numbers in parentheses represent mean and standard deviation obtained following histological verification of electrode locations. Trans-piriform electrodes were broken into those under the OFC and those under the NAc. See Figure 1 for recording sites across subjects. All units are in mm relative to bregma.

| Target | AP | ML | DV |
| --- | --- | --- | --- |
| Prelimbic (PL) | 3.0 (3.0±0.0) | 0.7 (0.5±0.3) | 3.2 (3.1±0.5) |
| Infralimbic (IL) | 3.0 (3.0±0.0) | 0.7 (0.5±0.2) | 4.4 (4.4±0.2) |
| Orbitofrontal cortex (OFC) | 3.0 (2.9±0.2) | 2.7 (2.7±0.3) | 5.4 (5.4±0.5) |
| Nucleus accumbens (NAc) | 1.5 (1.5±0.8) | 2.0 (2.5±0.9) | 7.6 (7.1±0.5) |
| Cingulate gyrus (CG) | 1.5 (1.7±0.5) | 0.8 (1.0±0.2) | 3.2 (2.6±0.3) |
| **Trans-Piriform** |  |  |  |
| Piriform - OFC | 3.0 (2.7±0.3) | 2.5 (2.9±0.1) | 6.9 (7.2±0.3) |
| Piriform - NAc | 1.5 (1±0.9) | 2.0 (3.0±1.0) | 8.1 (8.5±0.2) |

### Data acquisition and preprocessing

Local field potentials (LFPs) were acquired using a Digital Lynx data acquisition system with an HS-18MM preamplifier (Neuralynx), sampling LFP data at 2 kHz using a 1-400 Hz bandpass filter. Signals were referenced against ground (a skull screw above the ipsilateral cerebellum connected to system ground) in order to eliminate signal contamination from a common reference. Large-amplitude mechanical artifacts (>4 SD of the unfiltered data) and EMG activity (>3 SD from the mean of the signal envelope filtered between 200-500 Hz) were identified and removed from the LFP data.

### Naris occlusion

The main experimental manipulation in this study is the unilateral blockage of the nostrils (“naris occlusion”), alternating between ipsilateral (same side as LFP recording) and contralateral (opposite side). The naris closure protocol was identical to that described in Carmichael et al. (2017). Briefly, naris blockage tubes were constructed from PE90/100/110 tubing (BD Intramedic) by threading a human hair tied in a double overhand knot to a suture threaded through the inner wall of the tubing and then threading it back through the outer wall before protruding from the front of the tube. The hair and thread knot were then glued inside the tube and the protruding hair was cut to *∼*8 mm beyond the opening of the tubing, to allow for removal (Kucharski and Hall, 1987; Cummings et al., 1997; Carmichael et al., 2017). To insert the blockage tube, rats were placed under isoflurane anesthesia. The occlusion tube was first coated in sterilized Vaseline and then placed inside either nasal passage ipsi- or contralateral to the recording implant. Each subject underwent four recording sessions over the course of five days, with the third day acting as a break to ensure there was no swelling in the nasal passage due to repeated insertion and removal of tubes. Each recording session consisted of four 10-minute segments (Figure 4A): a non-occlusion baseline (“pre”), ipsilateral (“ipsi”) and contralateral (“contra”) naris occlusions (order counterbalanced across sessions) followed by another non-occlusion baseline (“post”). All segments were separated by >45 mins to minimize any effects of isoflurane anesthesia. Data was acquired while the rat rested on a cloth-covered flower pot, which is appropriate for analyzing gamma-band LFP oscillations because in anterior limbic structures such as the NAc gamma oscillations have been shown to be most prominent during resting periods rather than active running (Malhotra et al., 2015).

### Data analysis overview

Two complementary types of analysis were performed: (1) single-site analyses using power spectra, cross-frequency correlations, and number of detected gamma events; and (2) paired-site analyses using both non-directional (amplitude correlation and phase coherence) and directional (amplitude cross-correlation and phase slope) measures. All analyses, described in detail below, were performed using MATLAB 2014b (MathWorks) and can be reproduced using code and data available upon request.

### Spectral analysis

To determine the spectral content of the recorded LFPs, power-spectral densities (PSDs) were computed across each session segment (experimental condition) using Welch’s method (2048 sample Hanning window, with a 1024 sample overlap and a 4096 sample NFFT). For clarity, the 1/*f* trend was removed from the data by computing the PSD on the first derivative of the data (MATLAB `pwelch(diff(data))`, as per Carmichael et al. 2017), which we refer to as “whitened”. To quantify the amount of change in the power of a particular frequency range across experimental conditions, an area under the curve (AUC) measure was used. First, a curve was fit to the PSD from the “control” condition using the MATLAB `fit` function. A two-term exponential model was chosen based on visual inspection of the curve shape (MATLAB Curve Fitting Toolbox). The AUC between the PSD for each experimental condition and the control curve between the band of interest was then computed (see Figure 5A for an example). Spectrograms and cross-frequency correlations were computed for each session segment using the MATLAB function `spectrogram` (rectangular window of 512 samples with 25% overlap) for frequencies between 1-120 Hz.

### Gamma event detection and analysis

The goal of the event detection is to identify the characteristic bursts in which gamma oscillations tend to occur. Following Catanese et al. (2016) and Carmichael et al. (2017), we first obtained the amplitude envelopes by filtering the preprocessed LFP data for all segments in a recording session (low-gamma: 45–65 Hz; high-gamma: 70–90 Hz) using a 5^th^ order Chebyshev filter (ripple dB 0.5) with a zero-phase filter (MATLAB `filtfilt`) and taking the magnitude of the Hilbert-transformed signal. Next, a detection threshold was identified as the 95^th^ amplitude percentile taken from the “pre” and “post” data, converted to raw (*µ*V) thresholds, and applied to the full session data to yield a set of initial gamma event candidates. This two-step approach provides a consistent threshold in the face of changes in mean power both across and within segments. Candidate events were kept if they did not co-occur with high voltage spindles (*>*4 SD in mean amplitude envelope filtered between 7–11 Hz), had more than 4 oscillation cycles, and had a variance score (variance in amplitude of the peaks and troughs, divided by the mean amplitude of the peaks and troughs) less than 1.5.

### Amplitude cross-correlation

To determine the temporal relationship between gamma-band LFP oscillations in each pair of sites, we first used session-wide amplitude correlations computed across frequencies (3-100 Hz in 1 Hz steps). This non-directional measure reflects instantaneous (at time lag 0) temporal coordination between gamma oscillations across sites. Next, we determined the session-wide shift value (“lag”) with the maximum correlation for each frequency. Similar to the methods outlined in Adhikari et al. (2010a), the instantaneous amplitude of filtered input signals (pairs of recording sites) was shifted in time (*±*100 ms) to obtain the time lag for which the amplitude cross-correlation was largest.

In addition to session-wide amplitude cross-correlations, we also computed event-based amplitude cross-correlations by first filtering the entire session into either the low- or high-gamma bands and then restricting the data to detected gamma events (*±*100 ms of data on either end of the detection threshold). Events detected on either channel of a pair of sites were eligible for further analysis. However, because amplitude cross-correlations are only meaningful if the amplitudes of the two signals are related, we compared the observed peak cross-correlation to a distribution of surrogate cross-correlation peaks obtained by shuffling the phases of the Fourier transform of one of the signals (100 shuffles; see Catanese et al. 2016). If the peak cross-correlation of the actual gamma events did not exceed one standard deviation of the phase-shuffled distribution, then that event was excluded from further analysis.

### Coherence metrics

Phase coherence was extracted for each pair of recording channels across the entire recording session and within each detected gamma event. Session-wide coherence was computed using a 2048 sample Hanning window with 50% overlap (MATLAB `mscohere`, NFFT = 4096). Session-wide coherograms used the Chronux toolbox (http://chronux.org/, Mitra and Bokil 2007) with a 1 s moving window in 50 ms steps.

### Phase difference metrics

Phase coherence is a non-directional measure, so we also sought to estimate lead/lag relationships based on phase slopes (Nolte et al., 2008; Catanese et al., 2016). Briefly, by taking the phase differences between two signals across frequencies and computing the slope of the phase differences, the direction and magnitude of any lead/lag relationship can be determined. Unlike Granger causality (a common directionality measure), phase slopes are robust to independent noise sources applied to a common signal (Nolte et al., 2008). In our implementation, the phase difference between two channels was first computed for each frequency by taking the angle of the cross-spectral power density (MATLAB `cpsd` 256 sample Hamming window, 50% overlap, NFFT: 1024). Phase slopes across frequencies were estimated with circular regression applied to the phase differences using a 9-point window.

## Results

### Experiment 1: multisite recording of gamma oscillations with piriform inactivation

#### Gamma oscillations are highly synchronous across piriform proximal regions

We sought to characterize gamma-band LFP oscillations within and across a number of brain structures in the rat limbic system, and determine their relationship with piriform cortex activity. To this end, LFPs were recorded in up to five regions simultaneously (prelimbic: PL, infralimbic: IL, orbitofrontal cortex: OFC, nucleus accumbens: NAc, and cingulate gyrus: CG; Figure 1A) using chronically implanted electrodes while male rats (n = 7) rested on a terracotta pot covered in a towel. Consistent with earlier reports, OFC and NAc in particular showed prominent, co-occurring gamma-band LFP oscillations in both the low- (45-65 Hz) and high-gamma bands (70-90 Hz; see Figure 1B-C for examples). These gamma oscillations tended to occur in characteristic events, which we detected using a threshold-crossing procedure (indicated by the blue and green shaded areas in Figure 1B-C; see *Methods* for details). PL/IL and CG LFPs contained more subtle gamma oscillations, best visible when aligned with the more prominent OFC and NAc events (dashed vertical line in Figure 1B-C; note only PL data is shown in this example). Because OFC and NAc are anatomically more proximal to the piriform cortex compared to PL and CG, this pattern of gamma-band LFP amplitudes is suggestive of a volume-conducted piriform cortex source, whose amplitude decays with distance.

The traces in Figure 1B-C suggest that gamma oscillations across structures are temporally coordinated. As an initial step in characterizing this coordination, we computed the phase coherence and amplitude correlations across frequencies for each pair of sites. For each pair of sites, a peak in the low-gamma band in particular was apparent in both the phase coherence and amplitude correlation (Figure 2, see Figure S1 for all pairs). Coherence values in the low-gamma band ranged from 0.25 *±* 0.16 (OFC-CG; anatomically distal pair) to 0.64 *±* 0.28 (OFC-NAc; most coordinated and anatomically proximal to piriform) and low-gamma amplitude correlations ranged from 0.28 *±* 0.16 (OFC-CG) to 0.68 *±* 0.25 (OFC-NAc). High-gamma coherence showed a similar pattern: 0.25 *±* 0.19 (OFC-CG) to 0.57 *±* 0.18 (OFC-NAc) and amplitude correlations 0.27 *±* 0.18 (OFC-CG) to 0.61 *±* 0.13 (OFC-NAc; see Figure S1 for the full matrix of all values for all pairs of sites).

**Figure 2:**
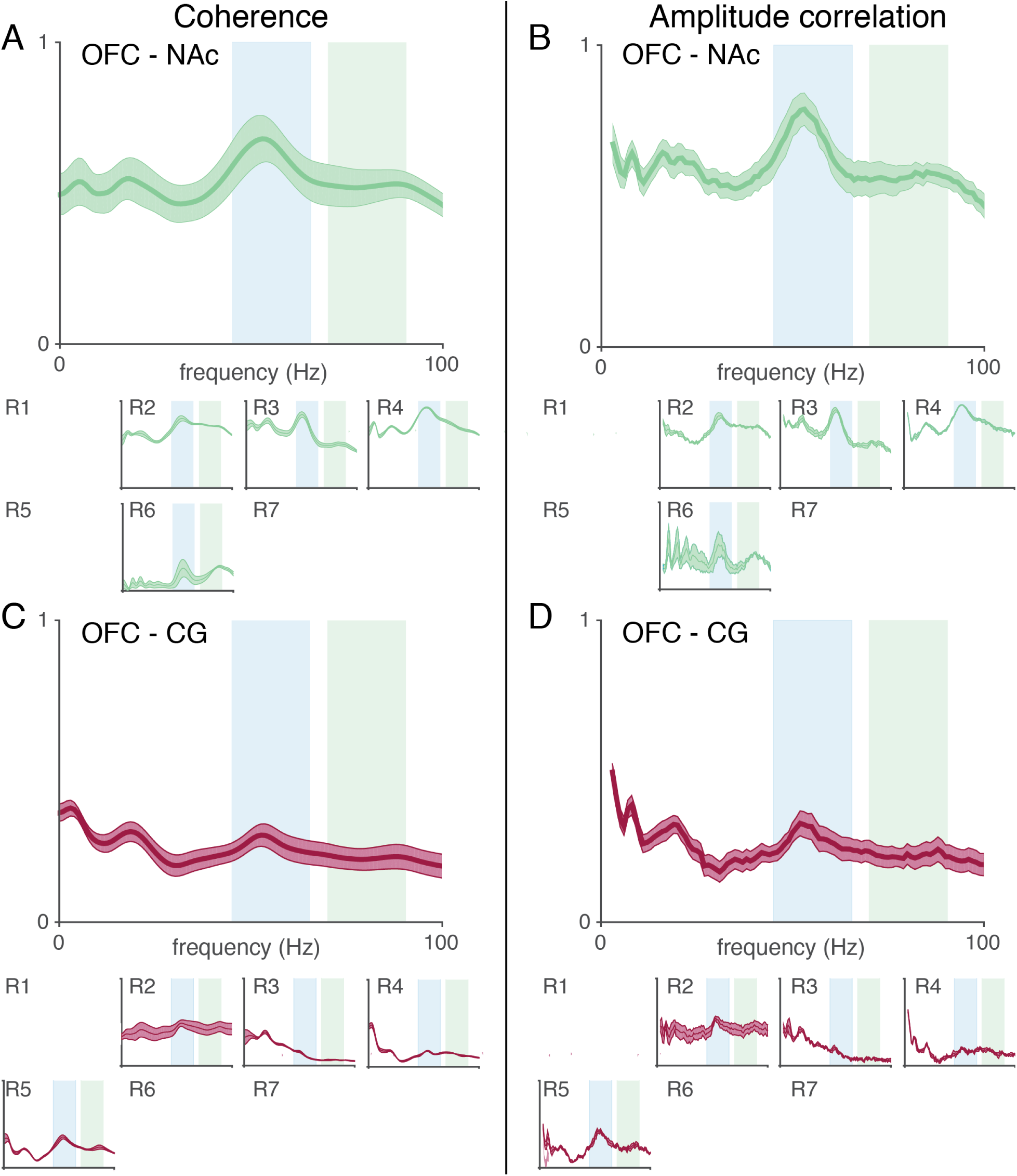
Pairs of areas proximal to piriform cortex, such as the OFC and NAc, show higher coherence and amplitude correlation compared to the more distal OFC-CG pair. Mean coherence (**A**) and amplitude correlation (**B**) across all subjects is elevated in the low-gamma band between the OFC and NAc (blue shades). This pattern is consistent across all subjects with electrodes in the OFC and NAc (lower). Coherence (**C**) and amplitude correlation (**D**) are low for the distal OFC-CG pair, with the exception of subject R5 which showed a moderate peak in the low-gamma band. Shaded areas represent SEM across sessions.

The above pattern of results indicates that sites close to piriform cortex, such as OFC-NAc, tended to be highly synchronous overall compared to pairs of sites where one site was distal to piriform cortex, such as OFC-CG (compare Figure 2A-B with Figure 2C-D). To test more formally whether distance from piriform cortex predicted coherence and amplitude cross-correlation, we used a linear mixed-effects model with average amplitude correlation or coherence in the low- or high-gamma bands as the dependent variable and distance from piriform cortex as the predictor. To account for repeated measures of electrodes in the same position across sessions for each subject, both session and subject were used as random effects. Anatomical distance from piriform cortex (defined as the maximum distance for each of the sites in a pair) significantly improved the model for low- and high-gamma amplitude correlation (low-gamma amplitude corr likelihood ratio: 52.41, p *<* 0.001; high-gamma amplitude corr likelihood ratio: 27.51, p *<* 0.001) and coherence (low-gamma coherence likelihood ratio: 43.93, p *<* 0.001; high-gamma coherence likelihood ratio: 22.41, p *<* 0.001). All models which included distance from piriform outperformed baseline models using only subject and session as predictors. This observation suggests that piriform cortex contributes to the degree of temporal coordination between limbic system areas.

Thus far these analyses have shown that gamma-band LFP oscillations are temporally coordinated across limbic system brain areas. However, coherence and amplitude correlations are non-directional measures and therefore cannot reveal any systematic lead or lag relationships between areas. To address this, we computed amplitude cross-correlations and phase slopes for each pair of sites, focusing on events with clear gamma power in at least one of the sites in a pair. Candidate events were detected by thresholding band-passed power at the 95^th^ percentile in low- and high-gamma bands respectively, and subjected to further selection criteria (see *Methods*). Identified gamma events were transient and highly synchronous as demonstrated by the high correlation in the envelope of the filtered signals in both the low- and high-gamma bands between OFC and NAc (low: 0.90 *±* 0.13; high: 0.87 *±* 0.14; Figure 3A). The amplitude cross-correlation was smaller for OFC-CG (low: 0.36 *±* 0.35; high: 0.44 *±* 0.40; Figure 3B and S2). The temporal offset with the highest amplitude correlation was zero for OFC-NAc (low-gamma: 0.00 ms, high-gamma: 0.00 ms), indicating a remarkable degree of temporal synchrony (Figure 3A). Phase slope, an alternative measure of lead/lag between two signals, confirmed the absence of any temporal offset across the gamma bands in the OFC-NAc pair (low-gamma: 0.09 ms, high-gamma: 0.56 ms; Figure 3C and S3).

**Figure 3:**
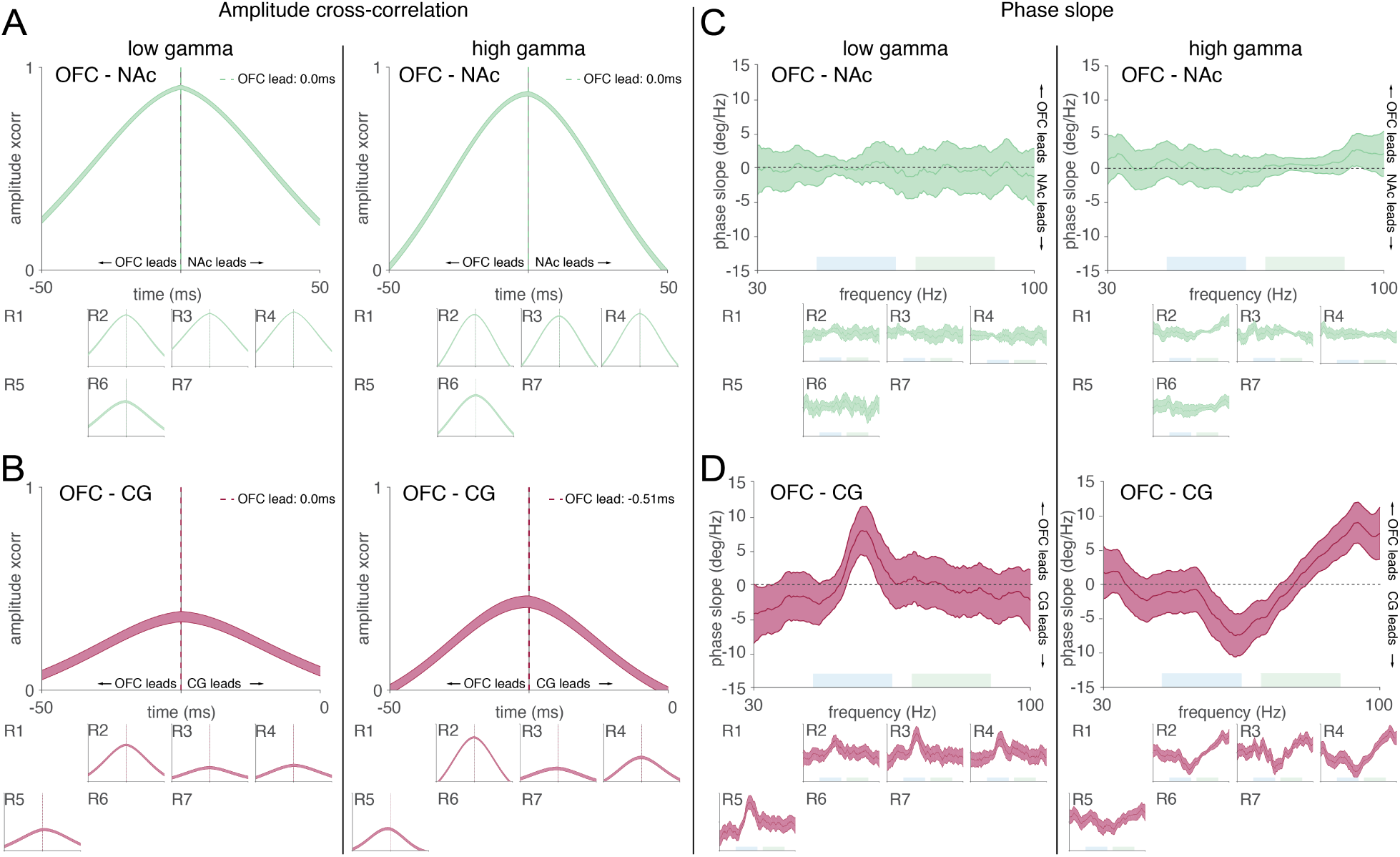
Gamma oscillations are highly synchronous across the limbic system, showing near-zero lag. Gamma event envelopes were highly correlated for both gamma bands in PC-proximal regions (**A**) and less correlated between distal pairs (**B**). Amplitude lead/lags were negligible for all electrode pairs, with the OFC showing a subtle lead over the CG in the high-gamma band. Individual subjects are shown below each cross-subject average plot. A minimum of 10 events was required for inclusion in the subject subplots (blank spaces represent subjects that did not have electrodes in both regions or failed to meet this criterion). Analysis of the phase slopes confirm a lack of lead/lag in the low- and high-gamma oscillations in the piriform-poximal regions (**C**). The OFC showed a small lead over the distal CG in both the low-gamma range for low-gamma events and a mild lead in higher frequencies (85-100 Hz) during high-gamma events (**D**). Shaded areas represent SEM across sessions.

**Figure 4:**
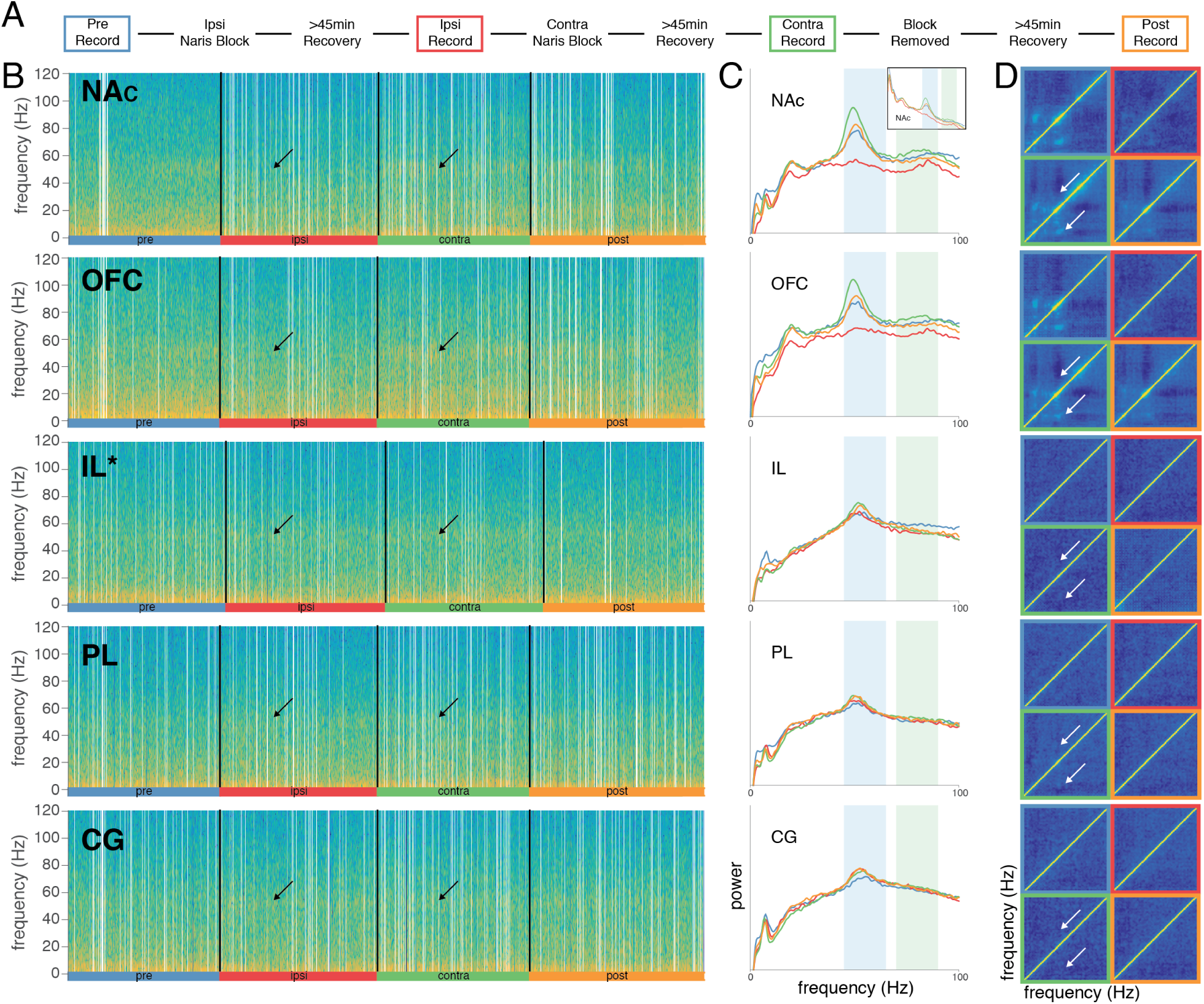
Blockage of the ipsilateral naris attenuates the gamma-band LFP power and disrupts cross-frequency correlations in regions proximal, but not distal, to the piriform cortex. (**A**) Naris occlusion protocol for a single session. “Ipsi” and “contra” phases were counterbalanced across sessions. (**B**) Spectrograms for an example session from a single subject (R3) across multiple sites, with the exception of “IL” which was not present in this subject and is from another subject (R4). During the “ipsi” phase there is a clear reduction in gamma-band power in the NAc, OFC, and to a lesser extent the PL and IL compared to the “contra” segment (arrows). Gamma power is still present during the “ipsi” segment for the CG. White lines represent artifacts. (**C**) Whitened power spectral densities (PSD) for the same sessions as in (**B**). Colors correspond to the four protocol phases in (**A**). Regions proximal to the piriform (NAc & OFC) show reduction in gamma power in the ipsilateral occlusion only, while distal regions show less pronounced changes. Inset: standard PSD for the same session which also shows the clear gamma band power reduction. (**D**) Cross-frequency autocorrelations reveal a common pattern in piriform-adjacent structures (NAc & OFC) with anti-correlation between low- and high-gamma, consistent with them not co-occurring, while showing high correlations between low-gamma and beta (15-30 Hz) bands and an anti-correlation between high-gamma and beta (white arrows). For piriform-adjacent sites these correlation patterns disappear with ipsilateral occlusion (red border).

**Figure 5:**
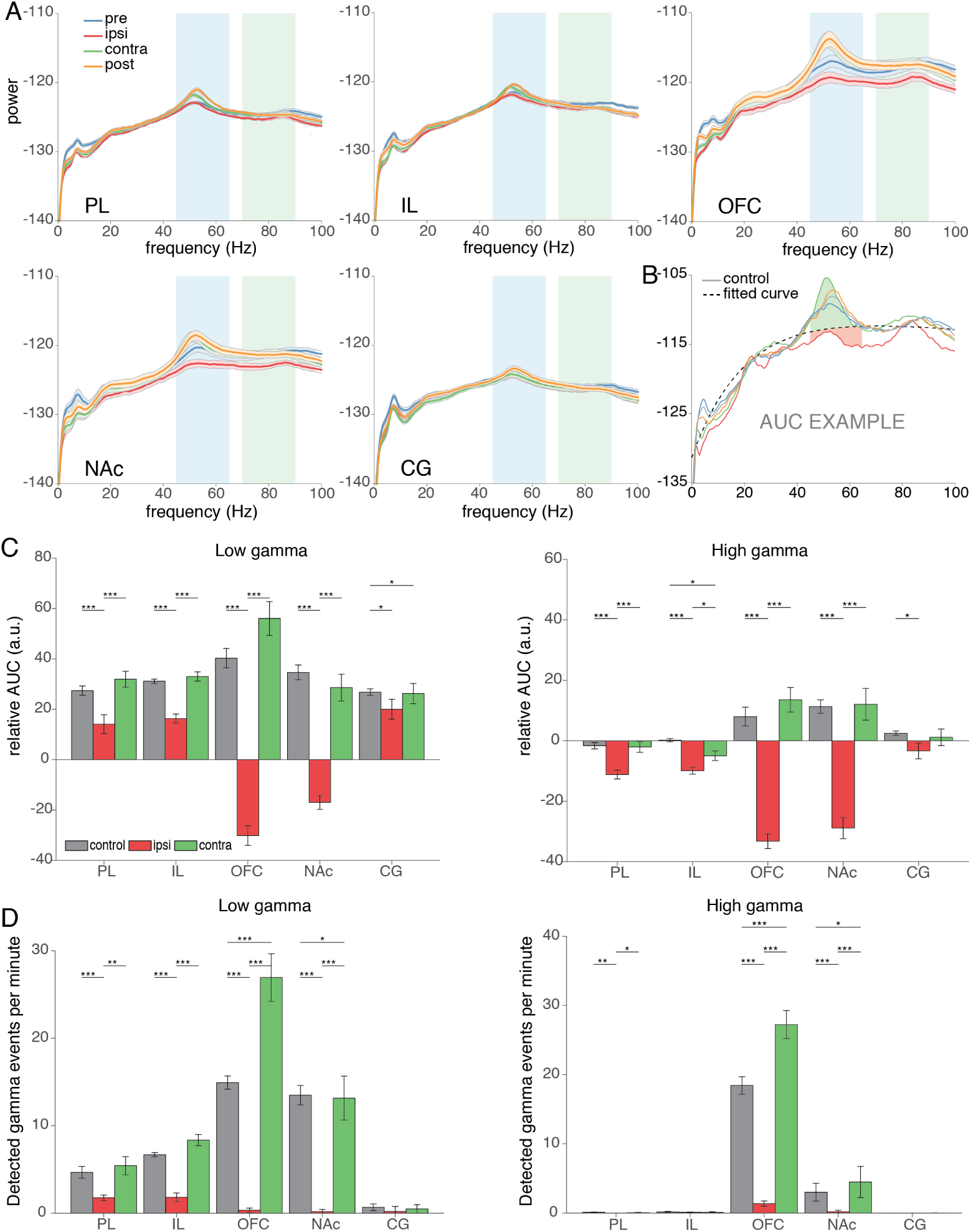
Unilateral naris occlusion decreases both gamma-band LFP power and the occurrence of detectable gamma events across regions. (**A**) Whitened power spectral densities averaged across all sessions and subjects for each recording site. Ipsilateral naris occlusion (red line) reduces the LFP power in the gamma bands with stark reductions in the OFC and NAc, and more subtle reductions in the PL, IL, and CG (shading represents the SEM). (**B**) To quantify the reduction in gamma power an exponential curve was fit to the “control” condition and the area under the curve (AUC) was calculated between each experimental condition relative to the fitted curve. (**C**) Summary of the relative AUC values for the low-gamma and high-gamma bands. (**D**) Ipsilateral nasal occlusion reduced the average number of detected gamma events per recording session compared to “control” or “contra” conditions. *p < 0.05, **p < 0.01, ***p < 0.001.

The OFC-CG pair showed little or no average OFC lead over the CG as measured with amplitude cross-correlation (low gamma: 0.00 ms, high gamma 0.51 ms; Figure 3B and S2). However, phase slopes displayed changes within each frequency band: for low-gamma, the OFC lead the CG (2.28 ms *±* 3.69) with a the greatest lead occurring between 55-60 Hz (7.03 ms *±* 0.79 ms). For high gamma events, the CG shows a lead over the OFC between *∼* 55-70 Hz (1.26 ms *±* 1.09 ms) while the OFC leads the CG between 70-90 Hz (1.03 ms *±* 3.61; Figure 3D and S3). These frequency-specific temporal relationships were highly consistent across subjects (see insets in Figure 3D) and illustrate the more precise view afforded by a frequency-resolved measure of directionality. Together with the session-wide coherence values these results illustrate the similarity of gamma signals in PC-proximal regions such as OFC-NAc, whose near-zero time offset indicates a common signal across the PC proximal regions. In contrast, gamma signals across areas that include a site more distal from PC such as OFC-CG, can show small deviations (on the order of a few milliseconds) from zero time offset.

#### Nasal occlusion disrupts gamma oscillations across the anterior limbic system

The tight temporal synchrony between areas shown above suggests that, consistent with Carmichael et al. (2017), PC may be the source of gamma oscillations and their coordination throughout the limbic system. To test this idea, we blocked olfactory and mechanical inputs to the nasal passage with a removable nose plug, applied either ipsi- or contralaterally to the recording electrodes. Such a blockage has previously been shown to disrupt gamma oscillations in the PC LFP (Zibrowski and Vanderwolf, 1997). The naris occlusion protocol (from Carmichael et al. 2017, outlined in Figure 4A) consisted of four recording segments: a pre-manipulation baseline (“pre”), occlusion of the nasal passage on the ipsilateral side of the recording electrodes (“ipsi”), occlusion of the contralateral nasal passage to the recording electrode “contra”, and a “post” recording at the end of the session. The order of the “ipsi” and “contra” segments were counterbalanced across days. For further analyses, we appended the “pre” and “post” segments to create a baseline “control” segment.

Figure 4B shows illustrative single-session spectrograms spanning all four recording segments. For all structures, power in the low-gamma band can be clearly seen as a horizontal band; however, for NAc and OFC in particular, power in this band was visibly reduced for the “ipsi” segment (red bar; arrows indicate comparison between “ipsi” and “contra”). This effect can also be seen in the corresponding power spectral densities (PSDs) for the same example session (Figure 4C): for NAc, OFC, and to some extent IL, the “ipsi” (red) and “contra” (green) PSDs show a clear attenuation of the peak in the low-gamma band, as well as a reduction in high-gamma power. For PL and CG, on the other hand, PSDs appeared very similar across “ipsi” and “contra” conditions. Next, we examined cross-frequency interactions, which are a hallmark of the NAc LFP (Sharott et al., 2009; van der Meer and Redish, 2009). As previously reported for NAc, low-gamma power was strongly correlated with beta power, while low- and high-gamma power were anticorrelated in both NAc and OFC, but not in the other areas (white arrows in Figure 4D in the “contra” segment indicated by the green border). During the “ipsi” condition these cross-frequency correlation patterns disappeared (red border). Thus, ipsilateral naris occlusion seems to affect gamma-band LFP oscillations not only in NAc, but also in OFC and to a lesser extent PL and IL; gamma oscillations CG appeared relatively unaffected by this manipulation.

To determine the generality of the above observation, we computed average PSDs across all subjects and sessions for each experimental condition (Figure 5A). The average PSDs confirm a systematic reduction in “ipsi” power compared to “contra” in both gamma bands for OFC and NAc in particular, and PL/IL to a lesser extent. In contrast, CG PSDs appeared similar across segments. To quantify the above observations, we fit a curve to the PSD for the “control” segment and calculated the area under the curve (AUC) for all segments relative to the fitted curve (example in Figure 5B). All regions except CG displayed a reduction in both low- and high-gamma power during the “ipsi” condition compared to either the “contra” or the “control” condition (Figure 5C). In CG, “ipsi” power was lower than control but not lower than contra for both low- and high-gamma; see Table 2 for the full set of linear mixed model comparisons).

**Table 2:** Summary of descriptive statistics and linear mixed effects model fits for gamma power and the rate of detected gamma events for naris occlusion conditions across sites. Power estimates are based on the area under the curve for the power spectral densities within the gamma bands relative to the “control” curve. Medians and SEM are reported in Figures 5C/D. *p < 0.05, **p < 0.01, ***p < 0.001.

| Area | Gamma power |  |  |  |  |  | Gamma event rate |  |  |  |  |  |
| --- | --- | --- | --- | --- | --- | --- | --- | --- | --- | --- | --- | --- |
|  | Naris condition | Low median | High median | Comparison | Low t-stat | High t-stat | Naris condition | Low median | High median | Comparison | Low t-stat | High t-stat |
| PL | Ipsi | 14.06±19.84 | -11.19±7.82 | Ipsi-Contra | -3.77 *** | -4.23*** | Ipsi | 1.76±1.70 | 0.00±0.06 | Ipsi-Contra | -3.56** | -2.50* |
|  | Contra | 31.97±16.81 | -2.05±9.40 | Ipsi-Control | -5.12*** | -6.19*** | Contra | 5.43±5.54 | 0.05±0.19 | Ipsi-Control | -4.78*** | -3.28** |
|  | Control | 27.40±9.73 | -1.64±5.58 | Contra-Control | 0.60 p = 0.55 | 0.77 p = 0.45 | Control | 4.67±3.56 | 0.10±0.16 | Contra-Control | -0.75 p = 0.46 | -0.13 p = 0.90 |
| IL | Ipsi | 16.32±9.58 | -9.95±5.88 | Ipsi-Contra | -4.71*** | -2.45* | Ipsi | 1.82±2.60 | 0.10±0.19 | Ipsi-Contra | -6.12*** | -0.99 p = 0.33 |
|  | Contra | 32.99±9.84 | -4.97±8.44 | Ipsi-Control | -5.03*** | -6.38*** | Contra | 8.36±3.34 | 0.10±0.62 | Ipsi-Control | -4.94*** | -1.46 p = 0.16 |
|  | Control | 31.13±4.55 | 0.18±2.82 | Contra-Control | 0.26 p = 0.79 | 2.56* | Control | 6.68±1.28 | 0.15±0.51 | Contra-Control | -1.76 p = 0.09 | 0.16 p = 0.88 |
| OFC | Ipsi | -30.16±20.43 | -33.23±13.01 | Ipsi-Contra | -10.54*** | -9.92*** | Ipsi | 0.34±1.39 | 1.36±2.04 | Ipsi-Contra | -8.12*** | -10.57*** |
|  | Contra | 56.08±35.50 | 13.55±21.51 | Ipsi-Control | -12.56*** | -9.92*** | Contra | 26.94±14.39 | 27.24±10.82 | Ipsi-Control | -18.46*** | -13.29*** |
|  | Control | 40.31±20.40 | 7.98±16.59 | Contra-Control | -1.73 p = 0.09 | -1.56 p = 0.13 | Control | 14.93±4.01 | 18.44±6.69 | Contra-Control | -3.75*** | -4.59*** |
| NAC | Ipsi | -16.95±14.90 | -28.86±18.62 | Ipsi-Contra | -10.35*** | -8.34*** | Ipsi | 0.18±1.42 | 0.20±1.04 | Ipsi-Contra | -6.87*** | -5.06*** |
|  | Contra | 28.62±28.05 | 12.06±27.68 | Ipsi-Control | -12.50*** | -11.13*** | Contra | 13.16±13.28 | 4.49±11.96 | Ipsi-Control | -11.53*** | -5.82*** |
|  | Control | 34.66±15.54 | 11.28±11.85 | Contra-Control | -1.12 p = 0.27 | -0.20 p = 0.84 | Control | 13.49±5.88 | 3.01±6.87 | Contra-Control | -2.55* | -2.54* |
| CG | Ipsi | 20.00±20.87 | -3.35±13.86 | Ipsi-Contra | -0.55 p = 0.58 | -0.79 p = 0.43 | Ipsi | 0.20±3.02 | 0.00±0.10 | Ipsi-Contra | 0.03 p = 0.98 | -0.56 p = 0.58 |
|  | Contra | 26.25±21.23 | 1.13±14.53 | Ipsi-Control | -2.78** | -2.47* | Contra | 0.49±2.56 | 0.00±0.17 | Ipsi-Control | -0.24 p = 0.81 | 0.58 p = 0.56 |
|  | Control | 26.81±6.83 | 2.49±3.97 | Contra-Control | 2.17* | 1.71 p = 0.10 | Control | 0.68±2.06 | 0.00±0.07 | Contra-Control | 0.34 p = 0.74 | -0.90 p = 0.38 |

Regions proximal to the PC appeared to display the largest reductions in both low- and high-gamma power during the “ipsi” condition, while distal regions showed more subtle reductions in gamma power. To determine if the anatomical distance from the PC for each electrode was a predictor of the effect of naris occlusion on reducing gamma-band power a linear mixed-effects model was used (MATLAB `fitlme`). Subject and session labels were included as random effects. Anatomical distance was a significant predictor of the contrast between ipsi and contra segments with piriform proximity showing the greatest reduction in gamma power (low-gamma: likelihood ratio: 59.91, p *<* 0.001; high-gamma: likelihood ratio: 69.82, p *<* 0.001). A similar model using the estimated anatomical distance from the olfactory bulb found it to be an ineffective predictor of the contrast between gamma power in ipsi and contra segments compared to distance form PC (low-gamma: likelihood ratio: 0.19, p = 0.66; high-gamma: likelihood ratio: 0.22, p = 0.64). These observations match what would be expected from volume conduction of LFP gamma oscillations from piriform cortex: in regions proximal to piriform cortex, volume conduction dominates and as a result the LFP is highly susceptible to naris occlusion. In contrast, in distal regions the contribution of volume conduction is smaller and the effect of naris occlusion is correspondingly minor.

To determine if the above reduction in gamma LFP power translates into different numbers of detected gamma events, we first used the ‘control’ condition to find an event threshold, and then applied this threshold to the ‘ipsi’ and ‘contra’ conditions to obtain event rates during these segments. For low-gamma, the event rate during ‘ipsi’ was significantly reduced compared to ‘contra’ for all regions except CG (CG p = 0.80, all other sites p *<* 0.01; see table 2 for full descriptors). For high-gamma, the “ispi” condition reduced the event rate in all regions except IL (IL p = 0.35, all other sites p *<* 0.05). All regions except CG showed a significant reduction in the low-gamma events rate during the “ipsi” condition compared to the “control” condition (CG p = 0.89, all other sites p *<* 0.001) while high-gamma saw a reduction in all regions except IL and CG (IL p = 0.16, CG p = 0.79, all other sites p *<* 0.001). Only the OFC (tstat: –3.69, p < 0.001) and NAc (tstat: –2.50, p < 0.05) displayed a significant increase in the occurrence of low-gamma events during the contralateral occlusion relative to the control condition. The average duration of the detected gamma events did not change as a result of the naris occlusion conditions for any of the sites. The reduction in detected gamma events in PC-adjacent regions mirrors the patterns of LFP power reduction (described above), and suggests that the gamma oscillations in NAc and OFC are not merely attenuated in power but are instead abolished entirely.

#### Nasal occlusion disrupts LFP synchrony across regions

Thus far, we have shown that gamma LFP oscillations in piriform-proximal regions are highly synchronous (session-wide and during gamma events), scale in amplitude with anatomical distance from the piriform cortex, and are susceptible to blockage of the nasal passage. These data suggest that the observed LFP synchrony across areas results from a common piriform cortex source. If this is true, then we would expect nasal occlusion to affect not only gamma oscillations in individual areas, but also their coordination across areas. To test this, we applied time-resolved measures of interregional connectivity (coherence and amplitude cross-correlation) to determine if the nasal occlusion shared a similar pattern of disruption to the reduction in gamma power reported above. The coherogram for the OFC-NAc pair shows a similar pattern to the spectrograms in Figure 4B with elevated coherence in the gamma band during the “pre”, “contra”, and “post” segments and a stark reduction in gamma coherence throughout the “ipsi” segment (Figure 6A). Momentary fluctuations in coherence can be seen in the “pre” and “post” segments of this example session, likely the result of changes in arousal level, but average coherence was systematically lower during “ipsi” compared to “contra” and control segments (Figure S1).

**Figure 6:**
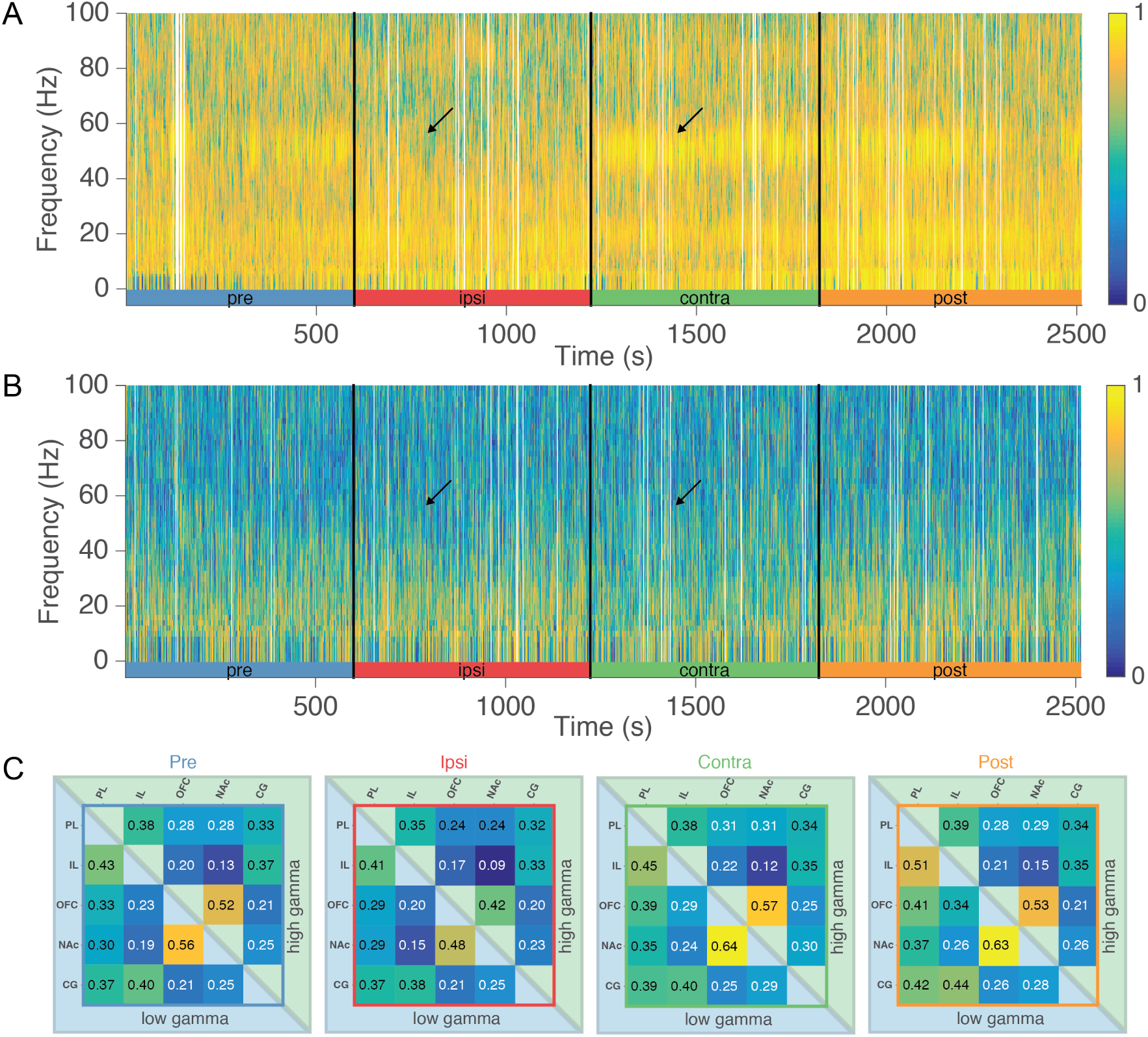
Unilateral naris occlusion decreases gamma-band coherence between regions proximal to the piriform cortex. (**A**) Representative coherogram across all four naris occlusion segments (pre, ipsi, contra, post) between the OFC and NAc for a single session. During the ‘ipsi’ segment there is a marked decrease in both low- and high-gamma (black arrows). White bars cover epochs with high amplitude artifacts. (**B**) Coherence between the OFC and the CG does not show any marked changes in coherence with the naris occlusion across any frequency bands. (**C**) Average low- and high-gamma band coherence across all subjects and sessions between all electrode pairs. During the four naris occlusion segments identified by the border of the matrices (’pre’: blue, ‘ipsi’: red, ‘contra’: green, ‘post’: orange). The lower triangle (blue) contains values in the low gamma band, while the upper triangle (green) contains high-gamma band data. For instance, sites that are proximal to one another display higher average coherence than distal sites. In addition, sites proximal to the piriform cortex display reduced coherence during the ‘ipsi’ segment relative to all other segments.

Piriform-distal electrode pairs (illustrated here by the OFC-CG pair) do not display elevated coherence in the gamma bands, nor does the nasal occlusion appear to greatly affect gamma-band coherence (Figure 6B). Piriform-proximal pairs of electrodes display increased session-wide coherence compared to more distal pairs (Figure 6C, see also Figure S1), with nasal occlusion reducing the coherence during that segment. Time-resolved amplitude cross-correlations mirrored the coherograms, suggesting that the changes in gamma power fluctuate together in time for piriform-proximal regions (Figure 7A), but not in piriform-distal pairs (Figure 7B). Nasal occlusion reduces the amplitude cross-correlation in the gamma band relative to the other segments (Figure 7C). These time-resolved measures of connectivity provide further support for a common volume-conducted gamma oscillation in the LFP across regions that are proximal to the piriform cortex.

**Figure 7:**
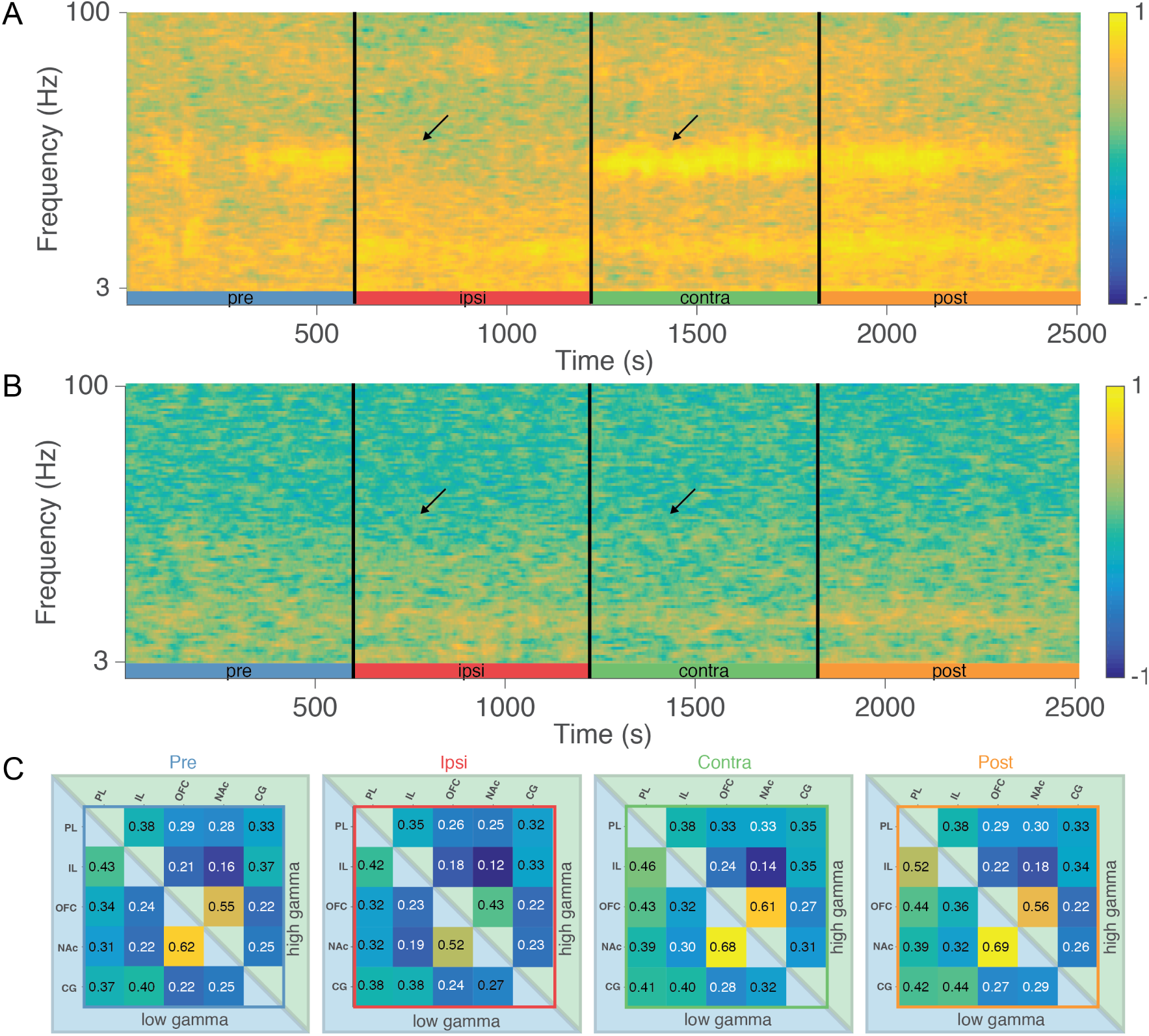
Naris occlusion reduces gamma-band amplitude cross correlation in proximal but not distal electrode pairs. (**A**) Representative amplitude cross-correlogram between the OFC and NAc. Naris occlusion only reduces amplitude cross-correlation in the gamma bands when the ipsilateral nasal passage is blocked. (**B**) Amplitude cross-correlogram between the OFC and the CG. The amplitude cross-correlation in the gamma bands is consistently low across all four segments, with no clear effect of the naris occlusions. (**C**) Summary of the amplitude cross-correlation values for each electrode par averaged across all subjects and sessions for low-gamma (blue, lower) and high-gamma (green, upper). The border of each matrix corresponds to the segment of the naris protocol in (**A**) & (**B**). Note the reduction in mean amplitude cross-correlation during the “ipsi” segment (red) compared to the other segments.

### Experiment 2: trans-piriform recording

Because all brain regions examined are anatomically located dorsal to the piriform cortex cell layer, volume conduction predicts that LFP phase differences across these regions are very close to zero (resulting from the near-instantaneous electrical propagation of the piriform LFP signal). In contrast, phase differences with sites ventral to the piriform cortex cell layer should display a 180 degree phase inversion, indicative of a sink/source pair located in the piriform cortex. To test this idea, we recorded simultaneously from sites located dorsal and ventral to the piriform cell layer (Figure 8A). If the ventral site was located in an area of piriform below the NAc or OFC, we refer to it as “Piri-NAc” and “Piri-OFC” respectively.

**Figure 8:**
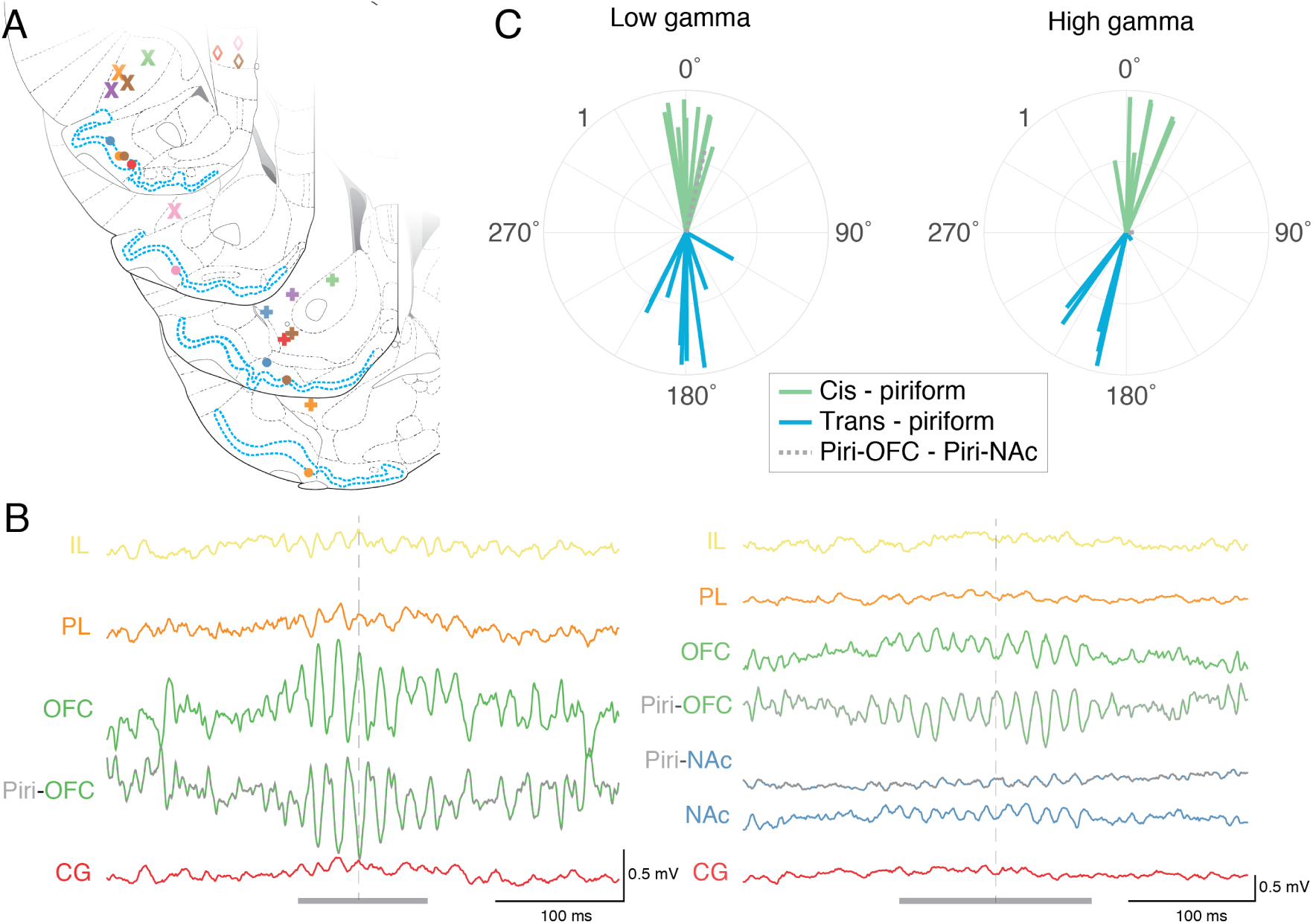
Recording across the piriform cortex reveals a consistent phase inversion in gamma band oscillations. (**A**) Histology for electrodes spanning the piriform cortex cell layer (highlighted in blue). (**B**) Example traces from low-gamma events (grey bar). A clear phase inversion can be seen between trans-piriform electrode pairs during the gamma events (dashed line). (**C**) Mean phase offsets for electrode pairs on the same side of the piriform cortex (”cis-piriform” in green) and those crossing the piriform cell layer (”trans-piriform” in blue) during the contralateral nasal occlusion. Cis-piriform electrodes displayed a consistent near-zero phase offset, while trans-piriform electrodes show a 180° offset. No differences emerge in the phase offsets patterns for either cis- or trans-piriform pairs between the contra and ipsi conditions (not shown).

Example traces recorded from either side of the PC cell layer show a clear phase inversion, both when recorded below the OFC (Piri-OFC, Figure 8B) and below the NAc (Piri-NAc, Figure 8B). Although gamma oscillations in PL, IL and CG were smaller than those in piriform-adjacent areas, as noted above, their phase was consistent with OFC/NAc, rather than inverted as was the case below the PC layer. To characterize these phase relationships across all recordings, we categorized each pair of recording sites either as being on the same side of the piriform cortex cell layer (i.e. located dorsal to it; “cis-piriform”) or as being on the opposide side (i.e. crossing the layer; “trans-piriform”). For example, OFC/PL and NAc/CG are cis-piriform pairs, whereas OFC/Piri-OFC and IL/Piri-NAc are trans-piriform pairs.

Using this categorization, we plotted the phase difference for each pair of recording sites as a line in a polar plot, such that its angle indicates the average phase difference, and its length indicates the variance (mean vector length across detected gamma events, Circular Statistics Toolbox, Berens 2009). Cis-piriform pairs consistently had phase differences near 0° for both low- and high-gamma events, as expected; in contrast, trans-piriform pairs had phase differences near 180°, indicative of a sink/source pair (Figure 8C). We noted a possible deviation from exactly 180° for high-gamma trans-piriform pairs; the reasons for this are currently unclear. Nevertheless, the clear phase reversal observed across the piriform cell layer at multiple locations demonstrates that the piriform cortex is the source of LFP gamma oscillations throughout the areas examined.

## Discussion

Inter-area synchrony in the neural activity of multiple brain areas is thought to reflect functional interactions, and perhaps even offer a mechanistic explanation for dynamic gain control (Fries, 2015). Local field potentials (LFPs) are often used as a proxy to measure synchrony, as found in the dynamic synchronization of limbic system LFPs across the nucleus accumbens, orbitofrontal and prefrontal cortex, amygdala and others (e.g. Gordon and Harris, 2015). Changes in limbic system synchrony correlate with specific behaviors and abnormal synchrony may be indicative of pathological states, motivating studies that investigate where limbic system LFPs are generated and what causes them to synchronize.

In this study, we have shown that gamma-band oscillations in the local field potential across the NAc, OFC, and PL/IL are highly similar, as indicated by their amplitude correlation and coherence. Both amplitude cross-correlations and phase slopes showed near-zero time lags across regions, indicating a high degree of temporal synchrony. These regions have in common that they are anatomically proximal to the piriform cortex (PC). We found that LFP gamma oscillations in these proximal regions were susceptible to ipsilateral nasal blockage, a manipulation known to abolish piriform gamma oscillations. Next, we identified the PC as the source of these common gamma oscillations because of the characteristic 180° phase reversal across its cell layer. Together, these results identify volume conduction from the piriform cortex as the main source of the dominant and highly synchronous gamma LFP oscillations seen throughout the anterior limbic system. These results inform our understanding of how gamma rhythmic activity in LFP and spiking is coordinated across limbic areas, and have implications for the interpretation of previous studies and future work. We will discuss these in turn below, starting with a working model of gamma-band synchrony in the limbic system.

### A working model of PC-based gamma coordination across anterior limbic regions

This study has provided several lines of evidence that, when taken together, demonstrate that the strikingly synchronous gamma-band oscillations in the LFP of multiple limbic system regions are due to volume conduction from a common source – the piriform cortex. However, this interpretation raises a potential conundrum: how can there be phase-locking of neurons to volume-conducted LFP oscillations, as has been found in each of the regions we examined (Berke, 2009; van der Meer and Redish, 2009; van Wingerden et al., 2010b; Kalenscher et al., 2010; Howe et al., 2011; Morra et al., 2012; Insel and Barnes, 2015)? We suggest that this question can be resolved by taking into account direct synaptic inputs from PC projection neurons, and/or other inputs correlated with PC activity, whose activity is gamma-phase locked. Thus, we propose that gamma-band spike-field locking in the anterior limbic system occurs because spike timing is inherited from an input that is also the source of the (non-local) field potential. This idea is summarized graphically in Figure 9.

**Figure 9:**
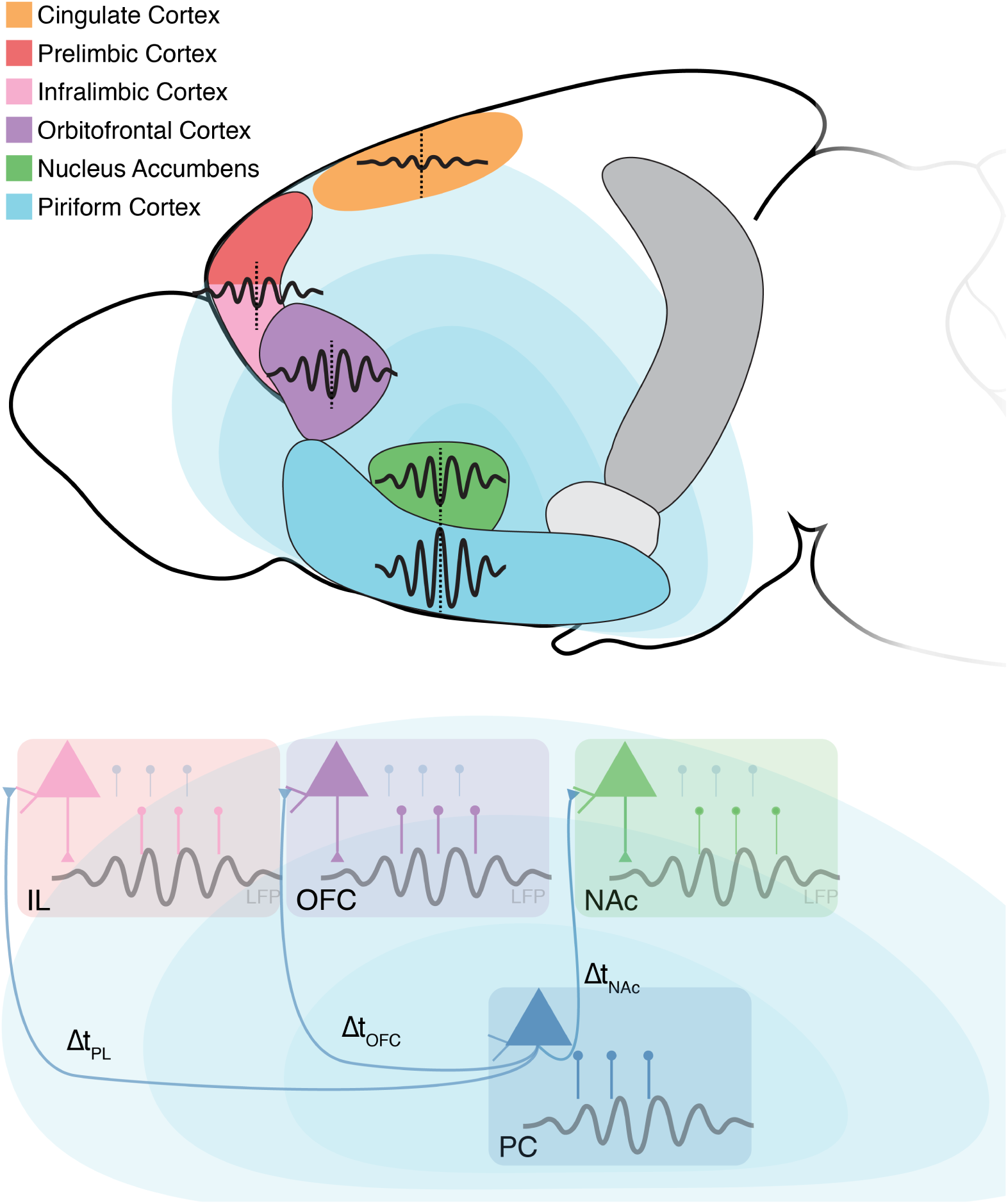
Schematic of a working model that explains both (1) common, highly synchronous gamma-band LFP oscillations (light blue) throughout the anterior limbic system, and (2) spike-field relationships. *Upper*: LFP gamma oscillations are generated in the PC (blue) and volume-conduct into nearby limbic regions, scaling in amplitude with distance from the PC origin. There is a phase reversal across the PC layers and a near-zero phase lag between regions (vertical dashed line). *Lower*: Phase-locked spiking to the common gamma signal is due to anatomical projections from PC, causing “inherited” gamma-band rhythmic spiking in connected regions. Faded blue lines in each of the limbic regions represent PC spikes. Note that local spiking (colored lines) is offset from the PC spikes by Δt, this results in local spiking that is still phase-locked to the PC gamma oscillation but at a later phase in the gamma LFP. Based on mouse brain sagittal by Jonas Töle, licensed under CC-BY-1.0.

This model of gamma synchrony through PC coordination requires anatomical connections (direct or indirect) between the PC and participating limbic regions. The OFC, NAc and IL receive direct synaptic inputs from the ipsilateral PC, and project back to PC mostly ipsilaterally (Hurley et al., 1991; Berendse et al., 1992; Brog et al., 1993; Datiche and Cattarelli, 1996; Haberly, 2001; Ekstrand et al., 2001; Illig, 2005). In contrast, the PL and CG cortices have few or no direct PC inputs (Hoover and Vertes, 2007). These anatomical connections would allow OFC, NAc and IL to inherit gamma-band rhythmic spiking from PC inputs as proposed above (Figure 9). In principle, this model does not strictly require a monosynaptic connection; for instance, motor efference copy or other respiration-related signals could also provide gamma-rhythmic inputs to these regions. For instance, a respiration-related/efference copy 4 Hz oscillation has recently been shown to entrain spiking and oscillatory events (including NAc and mPFC gamma) across the limbic system (Karalis and Sirota, 2018). However, such an indirect mechanism would be expected to maintain less precise spike-timing compared to a direct input. In sum, both monosynaptic projections and indirect efference-copy signals may contribute to common gamma LFP rhythmicity and spike-field locking across the anterior limbic system.

An important requirement for this working model to be correct is that gamma-band LFP power should fall off linearly with distance to the piriform cortex. Although we have not measured this gradient across the entire anterior brain with a laminar recording array, we have shown that the effects of nasal blockage on LFP gamma power scale with the distance from the PC (with the highest reduction in structures closest to the PC; Figures 4 and 5). In addition, in previous work, we have recorded across the ventral striatum using a grid of electrodes and found a gradient in gamma power that decreases with distance from the PC (Carmichael et al., 2017). The near-zero phase offsets between the limbic structures (Figure 8B) and across electrode grids in the vStr (Carmichael et al., 2017) adds further support to the volume conduction model, as local sources should produce differences in phase (which we did not find in this study, or in our previous work). Subtle irregularities in the phase offsets for certain regions may be the result of PC sections with pronounced curvature; in addition, there likely are some truly local components of the LFP even in non-laminar structures, as demonstrated by evoked potentials in slice and in vivo experiments (e.g. Pennartz and Kitai, 1994; Albertin et al. 2002 for the NAc). Such local contributions, expected to be small because of the non-laminar nature of the NAc, may account for the lead/lag times on the order of a millisecond found in this study and our previous work (Catanese et al., 2016; Carmichael et al., 2017).

Although we have argued that the PC is the source of common LFP gamma, gamma oscillations are highly correlated along the olfactory pathway from the OB to the PC (Kay and Freeman, 1998; Beshel et al., 2007; Mori et al., 2013; Frederick et al., 2016) which raises the question: which part of the olfactory pathway is being picked up at each site? Our reason for focusing on the piriform is two-fold: first, the strongest gamma oscillations were found in the OFC and NAc sites which are closer to the PC than the OB, and second, power gradients in the NAc have pointed to a piriform source (Berke, 2009; Carmichael et al., 2017). We also found that degree of naris-based gamma suppression also scales with anatomical distance from the PC rather than the OB (see *Results* text). The precise contributions of a volume-conducted signal from the OB compared to the PC could be determined through local inactivations or transections of the OB outputs, as in Parabucki and Lampl (2017).

A different potential limitation of this study is that we used only resting data, because running has been shown to reduce NAc/PC gamma power relative to rest in the NAc (van der Meer and Redish, 2009; Malhotra et al., 2015). It is nevertheless possible that a task component could lead to a locally generated gamma LFP independent of the PC. However, arguing against this possibility is first of all a direct comparison of task and rest gamma events, which turned out to be virtually identical (both volume-conducted) in the NAc (Carmichael et al., 2017). Second, a number of studies have recorded gamma LFPs during various tasks, finding highly synchronous gamma across pairs of regions examined (Beshel et al., 2007; Mori et al., 2013; Ponsel et al., 2017). Finally, we would expect that if local LFP sources were possible, resting state with its emphasis on internal dynamics rather than stimulus-driven activity would be more likely to reveal them.

Our model in which piriform cortex acts as a source of widespread gamma-band synchrony across the limbic system is consistent with a wider literature that highlights the relationship between respiration and LFP signals in multiple brain regions (Tort et al., 2018; Herrero et al., 2017; Heck et al., 2017), including correlations between anterior limbic gamma and respiration in the mPFC (Ponsel et al., 2017; Biskamp et al., 2017), OFC (Mori et al., 2013), as well as other cortical (Ito et al., 2014; Cavelli et al., 2018) and subcortical areas (Karalis and Sirota, 2008). However, our focus is specifically on understanding the source(s) of limbic system LFP oscillations and their relationship to local spiking.

### Implications of a common PC volume-conducted LFP gamma oscillation: caveats and reinterpretations

The data and associated working model presented here have two major implications. The first is a reinterpretation of previously reported behavioral correlates of gamma oscillations in the LFP in the anterior limbic system, and the second is a modification of targets for intervention. We discuss these in turn below and finish by outlining a few specific experimental predictions that provide further tests of our proposal.

Starting with the first, our results imply that gamma oscillations recorded in the PC-adjacent regions are more reflective of PC processing than they are of truly “local” activity. For instance, LFP gamma oscillations in a variety of limbic regions have been related to task events such as the presentation of odor cues and the delivery of rewards (van Wingerden et al., 2010b; Pennartz et al., 2011; van Wingerden et al., 2014; Cho et al., 2015; Kalenscher et al., 2010; van der Meer and Redish, 2009; Fujisawa and Buzsáki, 2011); we suggest that recordings from piriform cortex would show similar, perhaps even clearer versions of these task associations. Studies of gamma LFP changes following systemic administration of dopaminergic drugs or endocannabinoids (Morra et al., 2012; Berke, 2009; Goda et al., 2013) could reflect effects on piriform activity, a possibility that is further supported by the high density of dopamine receptors in the piriform cortex (Le Moine et al. 1990; Rocha et al. 1998; note similarities in NAc/OT DA projections in Ikemoto 2007). Similarly, altered LFP oscillations in animals that model aspects of human disease have been reported in all of the areas studied here (Tass et al., 2003; Greenberg et al., 2006, 2010; McCracken and Grace, 2007; Chamberlain et al., 2008; Bourne et al., 2012), suggesting that abnormalities in piriform cortex may be at least partly responsible.

Of course, widespread spike-field locking to gamma and other bands in the LFP (NAc: Berke et al. 2004; Berke 2005, 2009; Kalenscher et al. 2010; Morra et al. 2012; Malhotra et al. 2015; Catanese et al. 2016; van der Meer et al. 2019; Gmaz et al. 2019; OFC: van Wingerden et al. 2010a,b) means that even the volume-conducted piriform LFP contains at least some information about local activity. However, a volume-conducted LFP is a very indirect measure of local spiking; hypothetically, activity in say, the NAc may be completely suppressed through an experimental manipulation while leaving the piriform LFP intact. Even without experimental intervention, the efficacy of piriform inputs to the NAc may be modulated depending on task or behavioral state, making the volume-conducted LFP an even more tenuous measure of local activity compared to a “true” LFP (which itself isn’t straightforward to interpret, Berke 2005; Sciamanna and Wilson 2011; Schomburg et al. 2012; Wilson 2015; Pesaran et al. 2018).

An illustrative example of reinterpreting previous findings comes from ketamine-induced high-frequency oscillations (HFO) in the NAc. Hunt et al. (2006) had previously shown that systemic ketamine injection results in increased HFOs in the NAc. However, the same group found HFOs were also present in the OB and actually led the NAc HFOs in time (Hunt et al., 2019). Furthermore, inactivating the OB through nasal blockage or direct muscimol infusion attenuated NAc HFOs, implying an olfactory source. In addition to gamma and HFO oscillations, beta oscillations (*∼*15-25 Hz) are prominent in the olfactory system (Neville, 2003; Martin and Ravel, 2014; Kay et al., 2009) as well as the OFC and NAc (Berke 2009; McCracken and Grace 2009; Leventhal et al. 2012) suggesting that they too may have a common olfactory source (see Figures 4C and 5A, but see the uniform beta power in Howe et al. 2011 which does not change with electrode location).

It is possible that the influence of the common PC gamma rhythm extends to other regions beyond those examined in this study. The piriform cortex lies proximal to other limbic regions such as the amygdala, a non-laminar structure which also shows prominent gamma oscillations in the LFP (Collins et al., 2001; Bauer et al., 2007; Popescu et al., 2009; Sato et al., 2011, 2012; Stujenske et al., 2014; Likhtik and Paz, 2015). Thus, we suggest that (1) previously held interpretations of LFP oscillations in limbic structures take into account proximity to the piriform cortex, or other known generators of oscillations in the LFP, and (2) experimental diligence is applied when assessing the origin of LFPs (for examples and methods see: Sirota et al. 2008; Vinck et al. 2011; Bastos and Schoffelen 2016; Lalla et al. 2017; Esghaei et al. 2017; Torres et al. 2019; Feng et al. 2019).

The second implication of our working model follows straightforwardly from the first: if “local” field potentials in a number of limbic areas really are non-local, then any attempt to *change* the LFP should be targeted to its true, non-local source. This is an important consideration for studies that have used limbic system LFPs as biomarkers (e.g. impulsivity/binge eating: Wu et al. 2018; Doucette et al. 2018; Dwiel et al. 2019 and schizophrenia: Lodge et al. 2009). For instance, targeting the NAc with deep brain stimulation is unlikely to change its field potential, unless the stimulation were to antidromically affect piriform cortex inputs. Indeed, piriform cortex itself may turn out to be a DBS target: the human olfactory system has gained momentum as both an effective predictor in a range of neurological conditions (Doty, 2017) and as a target for DBS in epilepsy (Young et al., 2018) due to its extensive connectivity suggesting that impairments in this system could have larger implications for diagnostics and interventions.

Finally, our model of inherited piriform gamma rhythmicity in connected limbic regions makes several specific predictions. First, we would expect the inherited spiking in regions downstream from the PC to systematically phase-lock to later phases of the ongoing volume-conducted gamma oscillations than the PC neurons themselves (as shown in Figure 9). Second, if the downstream spike-field locking to PC gamma rhythms is the result of inherited spiking rhythmicity that coincides with the volume-conducted gamma LFP, then abolishing the projections from the PC would result in preserved gamma oscillations in the LFP in adjacent areas, but with a loss of gamma spike-field locking. Conversely, it may be possible to manipulate local spiking activity and/or synaptic currents, say by pharmacologically or pharmacogenetically inhibiting NAc activity, while leaving the field potential relatively unaffected.

More speculatively, it is tempting to wonder whether the common piriform cortex input, and the resulting inter-area synchronization across limbic brain structures, has implications for communication between these areas. In addressing this and related questions, it would be important to determine to what extent the limbic LFP indicates fluctuations in excitability – an idea that could be tested by measuring the magnitude or probability of a response to a fixed stimulus depending on LFP phase (Carmichael and van der Meer, 2019). These ideas show how even though the piriform source we have identified is cause for caution, it also offers ways forward in understanding how neural activity across the limbic system is coordinated.

## Acknowledgements

We thank Andrew Alvarenga for the design and machining of the four-tetrode drives, Jimmie Gmaz for technical support and useful comments on the manuscript, and Youki Tanaka for reagents. This work was supported by Dartmouth College (Dartmouth Fellowship to JEC, and start-up funds to MvdM).

## Supplemental Material

**Figure 10:**
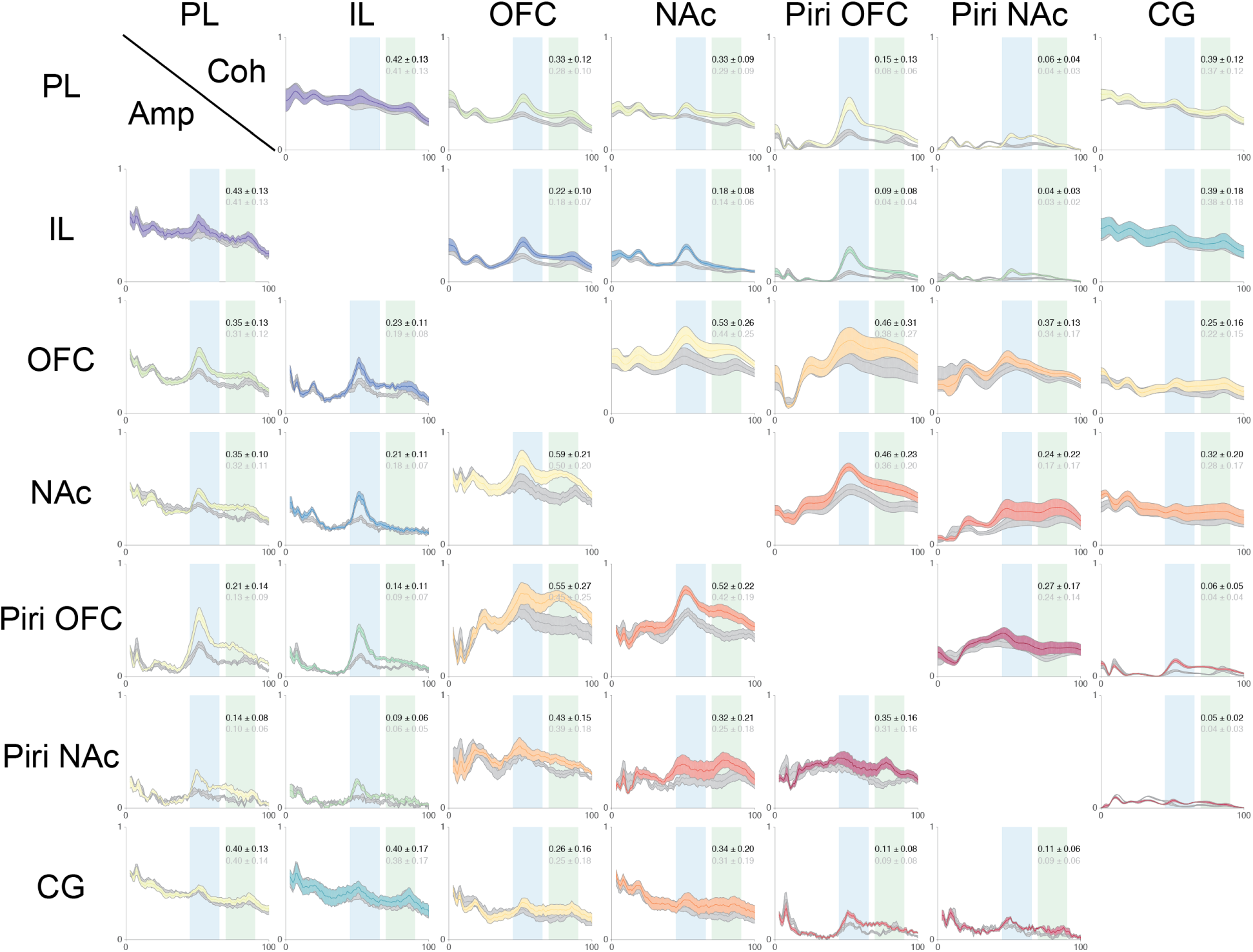
Mean coherence (upper) and amplitude cross-correlation (lower) across all sessions for each electrode pair. Solid colored lines represent the mean values for the contra condition across all sessions. Grey lines represent the mean values for the ipsi condition. Shaded areas represent one standard deviation. Nearly all electrode pairs show a prominent peak in the low gamma range (light blue rectangle) while electrode pairs with one or more sites proximal to the piriform cortex (OFC, NAc, Piri-OFC, Piri-NAc) show attenuation in the coherence or amplitude cross-correlation in the ipsilateral condition relative to the contralateral.

**Figure 11:**
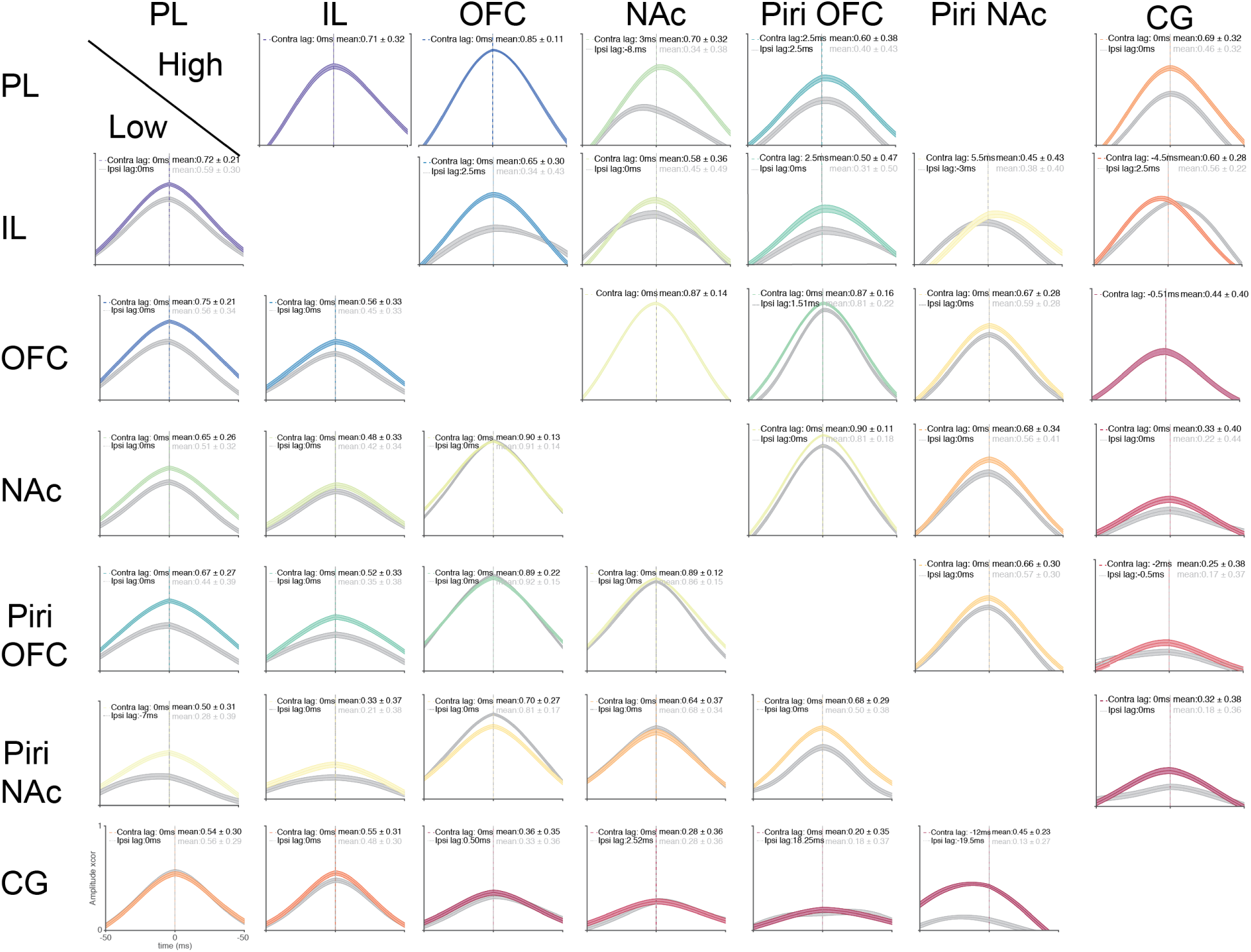
Event-based amplitude cross-correlation (low-gamma: lower, high-gamma: upper) across all sessions for each electrode pair. Solid colored lines represent the mean values for the contra condition across all sessions. Grey lines represent the mean values for the ipsi condition. Shaded areas represent one standard deviation. The maximum cross-correlation is highest for site pairs that are proximal to the piriform cortex and show a near-zero temporal delay, especially compared to distal electrode pairs. Ipsialteral nasal blockage reduces the amplitude cross-correlations.

**Figure 12:**
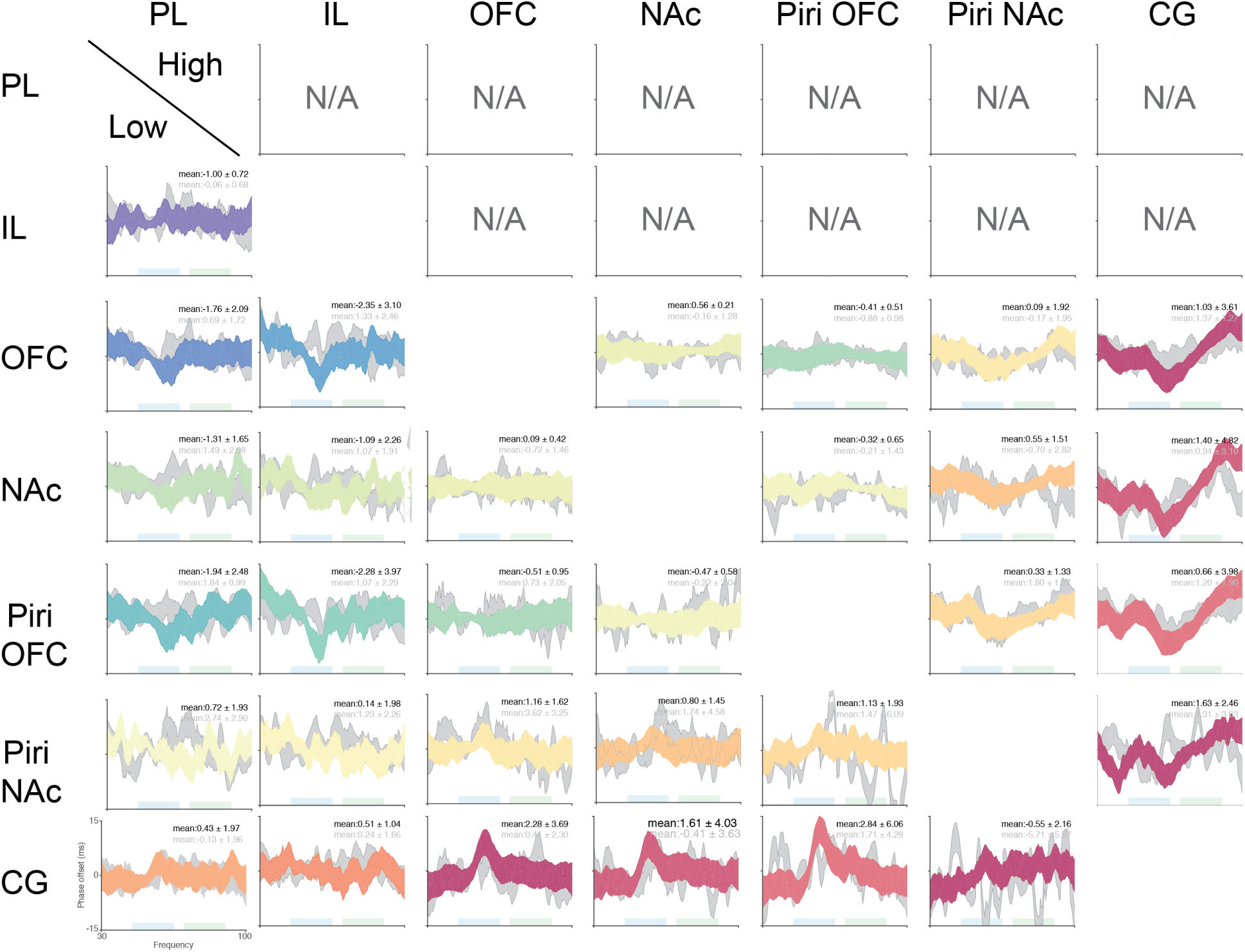
Event-based phase-slope values for low-gamma (lower) and high-gamma (upper) events across all sessions for each electrode pair. For consistency the low- and high-gamma event comparisons used the representations with positive values in the low-gamma phase slope representing a lead in the column site ID over the row site ID, while the high-gamma used the opposite with a positive value reprenting a lead in the row ID over the column ID (eg. in column 1 row 5 the PL lags behind the Piri-OFC in the low-gamma range). Solid colored lines represent the mean values for the contra condition across all sessions. Grey lines represent the mean values for the ipsi condition. Shaded areas represent one standard deviation. Sites with insufficient gamma events were excluded from analyses. PL and IL sites did not contain many high-gamma events which passed selection.

## References

1. Adhikari, A., Sigurdsson, T., Topiwala, M. A., and Gordon, J. A. (2010a). Cross-correlation of instantaneous amplitudes of field potential oscillations: a straightforward method to estimate the directionality and lag between brain areas. Journal of neuroscience methods, 191(2):191–200.

2. Adhikari, A., Topiwala, M. A., and Gordon, J. A. (2010b). Synchronized Activity between the Ventral Hippocampus and the Medial Prefrontal Cortex during Anxiety. Neuron, 65(2):257–269.

3. Bastos, A. M. and Schoffelen, J. M. (2016). A tutorial review of functional connectivity analysis methods and their interpretational pitfalls.

4. Bauer, E. P., Paz, R., and Paré, D. (2007). Gamma oscillations coordinate amygdalo-rhinal interactions during learning. The Journal of Neuroscience, 27(35):9369–79.

5. Berendse, H. W., Graaf, Y. G.-D., and Groenewegen, H. J. (1992). Topographical organization and relationship with ventral striatal compartments of prefrontal corticostriatal projections in the rat. The Journal of Comparative Neurology, 316(3):314–347.

6. Berens, P. (2009). CircStat: A MATLAB Toolbox for Circular Statistics. Journal of Statistical Software, 31(10):1–21.

7. Berke, J. D. (2005). Participation of Striatal Neurons in Large-Scale Oscillatory Networks. In The Basal Ganglia VIII, pages 25–36. Kluwer Academic Publishers, Boston.

8. Berke, J. D. (2009). Fast oscillations in cortical-striatal networks switch frequency following rewarding events and stimulant drugs. European Journal of Neuroscience, 30(5):848–859.

9. Berke, J. D., Okatan, M., Skurski, J., and Eichenbaum, H. B. (2004). Oscillatory entrainment of striatal neurons in freely moving rats. Neuron, 43(6):883–896.

10. Beshel, J., Kopell, N., and Kay, L. M. (2007). Olfactory bulb gamma oscillations are enhanced with task demands. Journal of Neuroscience, 27(31):8358–8365.

11. Biskamp, J., Bartos, M., and Sauer, J. F. (2017). Organization of prefrontal network activity by respiration-related oscillations. Scientific Reports.

12. Bonnefond, M., Kastner, S., and Jensen, O. (2017). Communication between Brain Areas Based on Nested Oscillations. eneuro, 4(2):ENEURO.0153–16.2017.

13. Bosman, C. A., Schoffelen, J. M., Brunet, N., Oostenveld, R., Bastos, A. M., Womelsdorf, T., Rubehn, B., Stieglitz, T., De Weerd, P., and Fries, P. (2012). Attentional Stimulus Selection through Selective Synchronization between Monkey Visual Areas. Neuron, 75(5):875–888.

14. Bourne, S. K., Eckhardt, C. A., Sheth, S. A., and Eskandar, E. N. (2012). Mechanisms of deep brain stimulation for obsessive compulsive disorder: effects upon cells and circuits. Frontiers in Integrative Neuroscience, 6:29.

15. Brog, J. S., Salyapongse, A., Deutch, A. Y., and Zahm, D. S. (1993). The patterns of afferent innervation of the core and shell in the ‘accumbens’ part of the rat ventral striatum: Immunohistochemical detection of retrogradely transported fluoro-gold. Journal of Comparative Neurology, 338(2):255–278.

16. Brown, P., Oliviero, A., Mazzone, P., Insola, A., Tonali, P., and Di Lazzaro, V. (2001). Dopamine dependency of oscillations between subthalamic nucleus and pallidum in Parkinson’s disease. The Journal of Neuroscience, 21(3):1033–8.

17. Carmichael, J. E., Gmaz, J. M., and van der Meer, M. A. A. (2017). Gamma Oscillations in the Rat Ventral Striatum Originate in the Piriform Cortex. The Journal of Neuroscience, 37(33):7962–7974.

18. Carmichael, J. E. and van der Meer, M. A. (2019). A physiological basis for communication through coherence in the rodent striatum. In 2019 Neuroscience Meeting Planner, page online, Chicago, IL. Society for Neuroscience.

19. Cassidy, M., Mazzone, P., Oliviero, A., Insola, A., Tonali, P., Lazzaro, V. D., and Brown, P. (2002). MovementâĂ Ř related changes in synchronization in the human basal ganglia. Brain, 125(6):1235–1246.

20. Catanese, J., Carmichael, J. E., and van der Meer, M. A. A. (2016). Low and high gamma oscillations deviate in opposite directions from zero-phase synchrony in the limbic corticostriatal loop. Journal of Neurophysiology, 116(March):jn.00914.2015.

21. Cavelli, M., Castro-Zaballa, S., Gonzalez, J., Rojas-Líbano, D., Rubido, N., Velásquez, N., and Torterolo, P. (2018). Nasal respiration entrains neocortical long-range gamma coherence during wakefulness. bioRxiv, page 430579.

22. Chamberlain, S. R., Menzies, L., Hampshire, A., Suckling, J., Fineberg, N. A., del Campo, N., Aitken, M., Craig, K., Owen, A. M., Bullmore, E. T., Robbins, T. W., and Sahakian, B. J. (2008). Orbitofrontal dysfunction in patients with obsessive-compulsive disorder and their unaffected relatives. Science (New York, N.Y.), 321(5887):421–2.

23. Cho, K., Hoch, R., Lee, A., Patel, T., Rubenstein, J., and Sohal, V. (2015). Gamma Rhythms Link Prefrontal Interneuron Dysfunction with Cognitive Inflexibility in Dlx5/6+/âĹŠ Mice. Neuron, 85(6):1332–1343.

24. Colgin, L. L. (2013). Mechanisms and Functions of Theta Rhythms. Annual Review of Neuroscience, 36(1):295–312.

25. Colgin, L. L., Denninger, T., Fyhn, M., Hafting, T., Bonnevie, T., Jensen, O., Moser, M.-B. B., and Moser, E. I. (2009). Frequency of gamma oscillations routes flow of information in the hippocampus. Nature, 462(7271):353–357.

26. Collins, D. R., Pelletier, J. G., and Paré, D. (2001). Slow and Fast (Gamma) Neuronal Oscillations in the Perirhinal Cortex and Lateral Amygdala. Journal of Neurophysiology, 85(4):1661–1672.

27. Concina, G., Cambiaghi, M., Renna, A., and Sacchetti, B. (2018). Coherent Activity between the Prelimbic and Auditory Cortex in the Slow-Gamma Band Underlies Fear Discrimination. The Journal of Neuroscience, 38(39):8313–8328.

28. Cummings, D. M., Henning, H. E., and Brunjes, P. C. (1997). Olfactory bulb recovery after early sensory deprivation. The Journal of Neuroscience, 17(19):7433–7440.

29. Datiche, F. and Cattarelli, M. (1996). Reciprocal and topographic connections between the piriform and prefrontal cortices in the rat: A tracing study using the B subunit of the cholera toxin. Brain Research Bulletin, 41(6):391–398.

30. DeCoteau, W. E., Thorn, C., Gibson, D. J., Courtemanche, R., Mitra, P., Kubota, Y., and Graybiel, A. M. (2007). Learning-related coordination of striatal and hippocampal theta rhythms during acquisition of a procedural maze task. Proceedings of the National Academy of Sciences, 104(13):5644–5649.

31. Dejean, C., Boraud, T., and Le Moine, C. (2013). Opiate dependence induces network state shifts in the limbic system. Neurobiology of Disease, 59:220–229.

32. Donnelly, N. A., Holtzman, T., Rich, P. D., Nevado-Holgado, A. J., Fernando, A. B. P., Van Dijck, G., Holzhammer, T., Paul, O., Ruther, P., Paulsen, O., Robbins, T. W., and Dalley, J. W. (2014). Oscillatory activity in the medial prefrontal cortex and nucleus accumbens correlates with impulsivity and reward outcome. PLoS ONE, 9(10):e111300.

33. Doty, R. L. (2017). Olfactory dysfunction in neurodegenerative diseases: is there a common pathological substrate? The Lancet Neurology, 16(6):478–488.

34. Doucette, W. T., Dwiel, L., Boyce, J. E., Simon, A. A., Khokhar, J. Y., and Green, A. I. (2018). Machine Learning Based Classification of Deep Brain Stimulation Outcomes in a Rat Model of Binge Eating Using Ventral Striatal Oscillations. Frontiers in Psychiatry, 9.

35. Dwiel, L. L., Khokhar, J. Y., Connerney, M. A., Green, A. I., and Doucette, W. T. (2019). Finding the balance between model complexity and performance: Using ventral striatal oscillations to classify feeding behavior in rats. PLoS Computational Biology.

36. Ekstrand, J. J., Domroese, M. E., Johnson, D. M. G., Feig, S. L., Knodel, S. M., Behan, M., and Haberly, L. B. (2001). A new subdivision of anterior piriform cortex and associated deep nucleus with novel features of interest for olfaction and epilepsy. Journal of Comparative Neurology, 434(3):289–307.

37. Esghaei, M., Daliri, M. R., and Treue, S. (2017). Local field potentials are induced by visually evoked spiking activity in macaque cortical area MT. Scientific Reports, 7(1).

38. Feng, F., Headley, D. B., Amir, A., Kanta, V., Chen, Z., Paré, D., and Nair, S. S. (2019). Gamma oscillations in the basolateral amygdala: Biophysical mechanisms and computational consequences. eNeuro, 6(1).

39. Frederick, D. E., Brown, A., Brim, E., Mehta, N., Vujovic, M., and Kay, L. M. (2016). Gamma and Beta Oscillations Define a Sequence of Neurocognitive Modes Present in Odor Processing. The Journal of Neuroscience, 36(29):7750–67.

40. Fries, P. (2005). A mechanism for cognitive dynamics: Neuronal communication through neuronal coherence. Trends in Cognitive Sciences, 9(10):474–480.

41. Fries, P. (2015). Rhythms for Cognition: Communication through Coherence. Neuron, 88(1):220–235.

42. Fujisawa, S. and Buzsáki, G. (2011). A 4 Hz Oscillation Adaptively Synchronizes Prefrontal, VTA, and Hippocampal Activities. Neuron, 72(1):153–165.

43. Gmaz, J. M., Carmichael, J. E., and van der Meer, M. A. (2019). Dynamic spike-field relationships in the rat nucleus accumbens. In 2019 Neuroscience Meeting Planner, page online, Chicago, IL. Society for Neuroscience.

44. Goda, S. A., Piasecka, J., Olszewski, M., Kasicki, S., and Hunt, M. J. (2013). Serotonergic hallucinogens differentially modify gamma and high frequency oscillations in the rat nucleus accumbens. Psychopharmacology.

45. Greenberg, B. D., Gabriels, L. A., Malone, D. A., Rezai, A. R., Friehs, G. M., Okun, M. S., Shapira, N. A., Foote, K. D., Cosyns, P. R., Kubu, C. S., Malloy, P. F., Salloway, S. P., Giftakis, J. E., Rise, M. T., Machado, A. G., Baker, K. B., Stypulkowski, P. H., Goodman, W. K., Rasmussen, S. A., and Nuttin, B. J. (2010). Deep brain stimulation of the ventral internal capsule/ventral striatum for obsessive-compulsive disorder: worldwide experience. Molecular Psychiatry, 15(1):64–79.

46. Greenberg, B. D., Malone, D. A., Friehs, G. M., Rezai, A. R., Kubu, C. S., Malloy, P. F., Salloway, S. P., Okun, M. S., Goodman, W. K., and Rasmussen, S. A. (2006). Three-Year Outcomes in Deep Brain Stimulation for Highly Resistant ObsessiveâĂ Ş - Compulsive Disorder. Neuropsychopharmacology, 31(11):2384–2393.

47. Gruber, A. J., Hussain, R. J., and O’Donnell, P. (2009). The nucleus accumbens: A switchboard for goal-directed behaviors. PLoS ONE, 4(4):e5062.

48. Haberly, L. B. (2001). Parallel-distributed Processing in Olfactory Cortex: New Insights from Morphological and Physiological Analysis of Neuronal Circuitry. Chemical Senses, 26(5):551–576.

49. Harris, A. Z. and Gordon, J. A. (2015). Long-Range Neural Synchrony in Behavior. Annual review of neuroscience, 38(March):171–194.

50. Heck, D. H., McAfee, S. S., Liu, Y., Babajani-Feremi, A., Rezaie, R., Freeman, W. J., Wheless, J. W., Papanicolaou, A. C., Ruszinkó, M., Sokolov, Y., and Kozma, R. (2017). Breathing as a Fundamental Rhythm of Brain Function. Frontiers in Neural Circuits, 10:115.

51. Herrero, J. L., Khuvis, S., Yeagle, E., Cerf, M., and Mehta, A. D. (2017). Breathing above the brainstem: Volitional control and attentional modulation in humans. Journal of Neurophysiology, page jn.00551.2017.

52. Hoover, W. B. and Vertes, R. P. (2007). Anatomical analysis of afferent projections to the medial prefrontal cortex in the rat. Brain Structure and Function, 212(2):149–179.

53. Howe, M. W., Atallah, H. E., McCool, a., Gibson, D. J., and Graybiel, a. M. (2011). Habit learning is associated with major shifts in frequencies of oscillatory activity and synchronized spike firing in striatum. Proceedings of the National Academy of Sciences, 108(40):16801–16806.

54. Hunt, M. J., Adams, N. E., Średniawa, W., Wójcik, D. K., Simon, A., Kasicki, S., and Whittington, M. A. (2019). The olfactory bulb is a source of high-frequency oscillations (130âĂŞ 180 Hz) associated with a subanesthetic dose of ketamine in rodents. Neuropsychopharmacology, 44(2).

55. Hunt, M. J., Raynaud, B., and Garcia, R. (2006). Ketamine Dose-Dependently Induces High-Frequency Oscillations in the Nucleus Accumbens in Freely Moving Rats. Biological Psychiatry, 60(11):1206–1214.

56. Hurley, K. M., Herbert, H., Moga, M. M., and Saper, C. B. (1991). Efferent projections of the infralimbic cortex of the rat. The Journal of Comparative Neurology, 308(2):249–276.

57. Igarashi, K. M., Lu, L., Colgin, L. L., Moser, M. B., and Moser, E. I. (2014). Coordination of entorhinal-hippocampal ensemble activity during associative learning. Nature.

58. Ikemoto, S. (2007). Dopamine reward circuitry: Two projection systems from the ventral midbrain to the nucleus accumbens-olfactory tubercle complex.

59. Illig, K. R. (2005). Projections from orbitofrontal cortex to anterior piriform cortex in the rat suggest a role in olfactory information processing. Journal of Comparative Neurology, 488(2):224–231.

60. Insel, N. and Barnes, C. A. (2015). Differential activation of fast-spiking and regular-firing neuron populations during movement and reward in the dorsal medial frontal cortex. Cerebral Cortex.

61. Ito, J., Roy, S., Liu, Y., Cao, Y., Fletcher, M., Lu, L., Boughter, J. D., Grün, S., and Heck, D. H. (2014). Whisker barrel cortex delta oscillations and gamma power in the awake mouse are linked to respiration. Nature Communications, 5(1):3572.

62. Jadi, M. P., Behrens, M. M., and Sejnowski, T. J. (2016). Abnormal Gamma Oscillations in N-Methyl-D-Aspartate Receptor Hypofunction Models of Schizophrenia. Biological Psychiatry, 79(9):716–726.

63. Jones, M. W. and Wilson, M. A. (2005). Theta Rhythms Coordinate HippocampalâĂŞ Prefrontal Interactions in a Spatial Memory Task. PLoS Biology, 3(12):e402.

64. Kajikawa, Y. and Schroeder, C. E. (2011). How local is the local field potential? Neuron, 72(5):847–858.

65. Kalenscher, T., Lansink, C. S., Lankelma, J. V., and Pennartz, C. M. A. (2010). Reward-Associated Gamma Oscillations in Ventral Striatum Are Regionally Differentiated and Modulate Local Firing Activity. Journal of Neurophysiology, 103(3):1658–1672.

66. Karalis, N., Dejean, C., Chaudun, F., Khoder, S., R Rozeske, R., Wurtz, H., Bagur, S., Benchenane, K., Sirota, A., Courtin, J., and Herry, C. (2016). 4-Hz oscillations synchronize prefrontal-amygdala circuits during fear behavior. Nature Neuroscience.

67. Karalis, N. and Sirota, A. (2018). Breathing coordinates limbic network dynamics underlying memory consolidation. bioRxiv, page 392530.

68. Kay, L. M., Beshel, J., Brea, J., Martin, C., Rojas-Líbano, D., and Kopell, N. (2009). Olfactory oscillations: the what, how and what for. Trends in Neurosciences, 32(4):207–214.

69. Kay, L. M. and Freeman, W. J. (1998). Bidirectional processing in the olfactory-limbic axis during olfactory behavior. Behavioral neuroscience, 112(3):541–53.

70. Kim, H., Ährlund-Richter, S., Wang, X., Deisseroth, K., and Carlén, M. (2016). Prefrontal Parvalbumin Neurons in Control of Attention. Cell, 164(1-2):208–218.

71. Kucharski, D. and Hall, W. G. (1987). New routes to early memories. Science, 238(4828):786–788.

72. Lalla, L., Rueda-Orozco, P., Jurado-Parras, M.-T., Brovelli, A., and Robbe, D. (2017). Local or Not Local: investigating the Nature of Striatal Theta Oscillations in Behaving Rats. Eneuro, 4(5):ENEURO.0128–17.2017.

73. Le Moine, C., Normand, E., Guitteny, A. F., Fouque, B., Teoule, R., and Bloch, B. (1990). Dopamine receptor gene expression by enkephalin neurons in rat forebrain. Proceedings of the National Academy of Sciences of the United States of America, 87(1):230–234.

74. Leventhal, D. K., Gage, G. J., Schmidt, R., Pettibone, J. R., Case, A. C., and Berke, J. D. (2012). Basal ganglia beta oscillations accompany cue utilization. Neuron, 73(3):523–536.

75. Likhtik, E. and Paz, R. (2015). Amygdala-prefrontal interactions in (mal)adaptive learning. Trends in neurosciences, 38(3):158–66.

76. Likhtik, E., Stujenske, J. M., Topiwala, M. A., Harris, A. Z., and Gordon, J. A. (2014). Prefrontal entrainment of amygdala activity signals safety in learned fear and innate anxiety. Nature neuroscience, 17(1):106–13.

77. Lodge, D. J., Behrens, M. M., and Grace, A. A. (2009). A loss of parvalbumin-containing interneurons is associated with diminished oscillatory activity in an animal model of schizophrenia. Journal of Neuroscience.

78. Logothetis, N. K., Kayser, C., and Oeltermann, A. (2007). In vivo measurement of cortical impedance spectrum in monkeys: implications for signal propagation. Neuron, 55(5):809–823.

79. Malhotra, S., Cross, R. W., Zhang, A., and Van Der Meer, M. A. A. (2015). Ventral striatal gamma oscillations are highly variable from trial to trial, and are dominated by behavioural state, and only weakly influenced by outcome value. European Journal of Neuroscience, 42(10):2818–2832.

80. Martin, C. and Ravel, N. (2014). Beta and gamma oscillatory activities associated with olfactory memory tasks: different rhythms for different functional networks? Frontiers in Behavioral Neuroscience, 8:218.

81. McCracken, C. B. and Grace, A. A. (2007). High-frequency deep brain stimulation of the nucleus accumbens region suppresses neuronal activity and selectively modulates afferent drive in rat orbitofrontal cortex in vivo. The Journal of Neuroscience, 27(46):12601–10.

82. McCracken, C. B. and Grace, A. A. (2009). Nucleus accumbens deep brain stimulation produces region-specific alterations in local field potential oscillations and evoked responses in vivo. J Neurosci, 29(16):5354–5363.

83. Mitra, P. and Bokil, H. (2007). Observed Brain Dynamics. Oxford University Press.

84. Moberly, A. H., Schreck, M., Bhattarai, J. P., Zweifel, L. S., Luo, W., and Ma, M. (2018). Olfactory inputs modulate respiration-related rhythmic activity in the prefrontal cortex and freezing behavior. Nature Communications, 9(1):1528.

85. Mori, K., Manabe, H., Narikiyo, K., and Onisawa, N. (2013). Olfactory consciousness and gamma oscillation couplings across the olfactory bulb, olfactory cortex, and orbitofrontal cortex. Frontiers in Psychology.

86. Morra, J. T., Glick, S. D., and Cheer, J. F. (2012). Cannabinoid receptors mediate methamphetamine induction of high frequency gamma oscillations in the nucleus accumbens. Neuropharmacology, 63(4):565–574.

87. Neville, K. R. (2003). Beta and Gamma Oscillations in the Olfactory System of the Urethane-Anesthetized Rat. Journal of Neurophysiology, 90(6):3921–3930.

88. Nolte, G., Ziehe, A., Krämer, N., Popescu, F., and Müller KRM, K.-R. (2008). Comparison of Granger Causality and Phase Slope Index. JMLR Workshop and Conference Proceedings.

89. Parabucki, A. and Lampl, I. (2017). Volume Conduction Coupling of Whisker-Evoked Cortical LFP in the Mouse Olfactory Bulb. Cell Reports, 21(4):919–925.

90. Pennartz, C. M., Van Wingerden, M., and Vinck, M. (2011). Population coding and neural rhythmicity in the orbitofrontal cortex. Annals of the New York Academy of Sciences, 1239(1):149–161.

91. Pesaran, B., Vinck, M., Einevoll, G. T., Sirota, A., Fries, P., Siegel, M., Truccolo, W., Schroeder, C. E., and Srinivasan, R. (2018). Investigating large-scale brain dynamics using field potential recordings: Analysis and interpretation. Nature Neuroscience.

92. Place, R., Farovik, A., Brockmann, M., and Eichenbaum, H. (2016). Bidirectional prefrontal-hippocampal interactions support context-guided memory. Nature Neuroscience, 19(8):992–994.

93. Ponsel, S., Draguhn, A., Ciatipis, M., Müller, C., Wolfenstetter, T., Yanovsky, Y., Brankačk, J., Tort, A. B. L., Zhong, W., and Jessberger, J. (2017). Selective entrainment of gamma subbands by different slow network oscillations. Proceedings of the National Academy of Sciences, 114(17):4519–4524.

94. Popescu, A. T., Popa, D., and Paré, D. (2009). Coherent gamma oscillations couple the amygdala and striatum during learning. Nature Neuroscience, 12(6):801–807.

95. Rocha, B. A., Fumagalli, F., Gainetdinov, R. R., Jones, S. R., Ator, R., Giros, B., Miller, G. W., and Caron, M. G. (1998). Cocaine self-administration in dopamine-transporter knockout mice. Nature Neuroscience, 1(2):132–137.

96. Sato, W., Kochiyama, T., Uono, S., Matsuda, K., Usui, K., Inoue, Y., and Toichi, M. (2011). Rapid Amygdala Gamma Oscillations in Response to Eye Gaze. PLoS ONE, 6(11):e28188.

97. Sato, W., Kochiyama, T., Uono, S., Matsuda, K., Usui, K., Inoue, Y., and Toichi, M. (2012). Temporal Profile of Amygdala Gamma Oscillations in Response to Faces. Journal of Cognitive Neuroscience, 24(6):1420–1433.

98. Schomburg, E. W., Anastassiou, C. A., Buzsáki, G., and Koch, C. (2012). The spiking component of oscillatory extracellular potentials in the rat hippocampus. Journal of Neuroscience.

99. Sciamanna, G. and Wilson, C. J. (2011). The ionic mechanism of gamma resonance in rat striatal fast-spiking neurons. Journal of Neurophysiology, 106(6):2936–2949.

100. Sharott, A., Moll, C. K. E., Engler, G., Denker, M., Grun, S., and Engel, A. K. (2009). Different Subtypes of Striatal Neurons Are Selectively Modulated by Cortical Oscillations. Journal of Neuroscience, 29(14):4571–4585.

101. Shin, H., Law, R., Tsutsui, S., Moore, C. I., and Jones, S. R. (2017). The rate of transient beta frequency events predicts behavior across tasks and species. eLife, 6.

102. Sigurdsson, T., Stark, K. L., Karayiorgou, M., Gogos, J. A., and Gordon, J. A. (2010). Impaired hippocampal-prefrontal synchrony in a genetic mouse model of schizophrenia. Nature.

103. Sirota, A., Montgomery, S., Fujisawa, S., Isomura, Y., Zugaro, M., and Buzsáki, G. (2008). Entrainment of Neocortical Neurons and Gamma Oscillations by the Hippocampal Theta Rhythm. Neuron, 60(4):683–697.

104. Sohal, V. S., Zhang, F., Yizhar, O., and Deisseroth, K. (2009). Parvalbumin neurons and gamma rhythms enhance cortical circuit performance. Nature, 459(7247):698–702.

105. Stujenske, J. M. M., Likhtik, E., Topiwala, M. A. A., and Gordon, J. A. A. (2014). Fear and Safety Engage Competing Patterns of Theta-Gamma Coupling in the Basolateral Amygdala. Neuron, 83(4):919–933.

106. Tass, P. A., Klosterkötter, J., Schneider, F., Lenartz, D., Koulousakis, A., and Sturm, V. (2003). Obsessive-Compulsive Disorder: Development of Demand-Controlled Deep Brain Stimulation with Methods from Stochastic Phase Resetting. Neuropsychopharmacology, 28(S1):S27–S34.

107. Torres, D., Makarova, J., Ortuño, T., Benito, N., Makarov, V. A., and Herreras, O. (2019). Local and Volume-Conducted Contributions to Cortical Field Potentials. Cerebral Cortex.

108. Tort, A. B. L., Brankačk, J., and Draguhn, A. (2018). Respiration-Entrained Brain Rhythms Are Global but Often Overlooked. Trends in Neurosciences.

109. Uhlhaas, P. J. and Singer, W. (2006). Neural Synchrony in Brain Disorders: Relevance for Cognitive Dysfunctions and Pathophysiology. Neuron, 52(1):155–168.

110. van der Meer, M. A. A., Gmaz, J. M., and Carmichael, J. E. (2019). A comprehensive characterization of rhythmic spiking activity in the rat ventral striatum. bioRxiv.

111. van der Meer, M. A. A. and Redish, A. D. (2009). Low and high gamma oscillations in rat ventral striatum have distinct relationships to behavior, reward, and spiking activity on a learned spatial decision task. Frontiers in Integrative Neuroscience, 3:9.

112. van Wingerden, M., van der Meij, R., Kalenscher, T., Maris, E., and Pennartz, C. M. A. (2014). Phase-Amplitude Coupling in Rat Orbitofrontal Cortex Discriminates between Correct and Incorrect Decisions during Associative Learning. J Neurosci, 34(2):493–505.

113. van Wingerden, M., Vinck, M., Lankelma, J., and Pennartz, C. M. (2010a). Theta-band phase locking of orbitofrontal neurons during reward expectancy. Journal of Neuroscience.

114. van Wingerden, M., Vinck, M., Lankelma, J. V., and Pennartz, C. M. A. (2010b). Learning-Associated Gamma-Band Phase-Locking of Action-Outcome Selective Neurons in Orbitofrontal Cortex. Journal of Neuroscience, 30(30):10025–10038.

115. Vinck, M., Oostenveld, R., Van Wingerden, M., Battaglia, F., and Pennartz, C. M. A. (2011). An improved index of phase-synchronization for electrophysiological data in the presence of volume-conduction, noise and sample-size bias. NeuroImage, 55(4):1548–1565.

116. Wilson, C. J. (2015). Oscillators and Oscillations in the Basal Ganglia.

117. Womelsdorf, T., Valiante, T. A., Sahin, N. T., Miller, K. J., and Tiesinga, P. (2014). Dynamic circuit motifs underlying rhythmic gain control, gating and integration. Nature Neuroscience, 17(8):1031–1039.

118. Wu, H., Miller, K. J., Blumenfeld, Z., Williams, N. R., Ravikumar, V. K., Lee, K. E., Kakusa, B., Sacchet, M. D., Wintermark, M., Christoffel, D. J., Rutt, B. K., Bronte-Stewart, H., Knutson, B., Malenka, R. C., and Halpern, C. H. (2018). Closing the loop on impulsivity via nucleus accumbens delta-band activity in mice and man. Proceedings of the National Academy of Sciences of the United States of America, 115(1):192–197.

119. Young, J. C., Vaughan, D. N., Paolini, A. G., and Jackson, G. D. (2018). Electrical stimulation of the piriform cortex for the treatment of epilepsy: A review of the supporting evidence. Epilepsy & Behavior, 88:152–161.

120. Zibrowski, E. M. and Vanderwolf, C. H. (1997). Oscillatory fast wave activity in the rat pyriform cortex: Relations to olfaction and behavior. Brain Research, 766(1-2):39–49.

